# Scalable gradients enable Hamiltonian Monte Carlo sampling for phylodynamic inference under episodic birth-death-sampling models

**DOI:** 10.1101/2023.10.31.564882

**Authors:** Yucai Shao, Andrew F. Magee, Tetyana I. Vasylyeva, Marc A. Suchard

## Abstract

Birth-death models play a key role in phylodynamic analysis for their interpretation in terms of key epidemiological parameters. In particular, models with piecewise-constant rates varying at different epochs in time, to which we refer as episodic birthdeath-sampling (EBDS) models, are valuable for their reflection of changing transmission dynamics over time. A challenge, however, that persists with current time-varying model inference procedures is their lack of computational efficiency. This limitation hinders the full utilization of these models in large-scale phylodynamic analyses, especially when dealing with high-dimensional parameter vectors that exhibit strong correlations. We present here a linear-time algorithm to compute the gradient of the birth-death model sampling density with respect to all time-varying parameters, and we implement this algorithm within a gradient-based Hamiltonian Monte Carlo (HMC) sampler to alleviate the computational burden of conducting inference under a wide variety of structures of, as well as priors for, EBDS processes. We assess this approach using three different real world data examples, including the HIV epidemic in Odesa, Ukraine, seasonal influenza A/H3N2 virus dynamics in New York state, America, and Ebola outbreak in West Africa. HMC sampling exhibits a substantial efficiency boost, delivering a 10to 200-fold increase in minimum effective sample size per unit-time, in comparison to a Metropolis-Hastings-based approach. Additionally, we show the robustness of our implementation in both allowing for flexible prior choices and in modeling the transmission dynamics of various pathogens by accurately capturing the changing trend of viral effective reproductive number.

## 1 Introduction

Phylodynamic models constitute a sophisticated toolset employed to decipher the complex interplay between epidemiological and evolutionary processes, providing valuable insights into population dynamics (Lau et al. 2019). In this paper, our primary emphasis is directed toward the inference of epidemiological dynamics, rather than estimation of the underlying phylogeny through sequence analysis. Specifically, we start with a sample of molecular sequences, which can be used to reconstruct the evolutionary relationships between organisms, often viral pathogens, and yield inference on dynamics of the larger pathogen population over time while relegating the phylogeny the status of a nuisance parameter. To provide this link, a vital component of phylodynamic analysis is the use of birth-death models, which belong to an important subclass of continuous-time Markov chains (CTMCs). We use birth-death models to define the probability distribution on time-calibrated phylogenies for reflecting the fluctuations of the population size (MacPherson et al. 2022). In this context, birth-death models posit three major types of events: birth, which refers to the creation of new lineages through pathogen transmission between hosts; death, which represents host death/recovery or other removal from the studied population, and sampling, which means the collection of a sequence derived from the pathogen in a single infected host and included in the data set under analysis (Crawford 2012).

The past few decades have delivered a wide range of birth-death models. These span from a simple, constant-over-time formulation (Yang & Rannala 1997) to models that allow both birth and death rates to vary over time (Stadler et al. 2013, Höhna 2014). Further extensions incorporate additional processes, both statistical and biological, such as the collection of samples in continuous time (Stadler 2010), migration (Barido-Sottani et al. 2020), or the dependency of rates of birth and death on key biological traits (Maddison et al. 2007, FitzJohn 2010, 2012). One powerful variant, the episodic birth-death-sampling (EBDS) model (Lambert & Stadler 2013, Stadler et al. 2013, Gavryushkina et al. 2014, Du Plessis 2016) permits birth, death, and sampling rates to change in discrete epochs throughout time to capture more complicated population dynamics. Recent inference based on EBDS models has found its way already into many applications, especially on the understanding of the spread of infectious disease (Novitsky et al. 2015, Vasylyeva et al. 2020, Minosse et al. 2021).

With increasingly rich and complex molecular sequence datasets across fields, improving the scalability of inference under EBDS models remains challenging both in terms of the number of sequences and the number of epochs. The most commonly employed inference methods based on Markov chain Monte Carlo (MCMC) (Hastings 1970, Morlon et al. 2011) use random-walk transition kernels generally to propose new parameter values in a blind fashion. Consequently, they lead to many birth-death model likelihood evaluations and slow exploration across the state space, especially for high-dimensional problems. The potentially complex correlation structure between epoch parameters can further exacerbate inference. This is where gradient-based sampling methods, such as Hamiltonian Monte Carlo (HMC) (Duane et al. 1987, Neal et al. 2011), are expected to shine. HMC has recently become very popular as a MCMC algorithm that overcomes many of the limitations of random-walk Metropolis-Hasting (MH) methods. Instead of making random proposals, HMC exploits the gradient of the log posterior with respect to (wrt) its model parameters to propose new states that are likely to be accepted and are far from the current state. Since HMC can make large moves in the state space while still maintaining a high acceptance rate, it can lead to faster convergence and better mixing than MH approaches, if one can efficiently evaluate not only the log posterior (up to a constant) but also its gradient. Successful implementation of HMC transition kernels has proved fruitful in terms of boosting sampling performance in other phylogenetic inference frameworks, including for different clock models (which describe how rates of molecular evolution vary among different organisms over time, Ji et al. 2020, Fisher et al. 2021), divergence times (the internal-node heights of phylogenies, Ji et al. 2021) and non-parametric coalescent models (which fall into another category of phylodynamic models assuming effective population size as a piecewise-constant form of time, Baele et al. 2020).

In this paper, we incorporate gradient-based sampling methods into phylodynamic analysis based on EBDS models, thereby enabling scalable inference within this framework. First, we refactor the EBDS (log) likelihood to show explicitly that the computational complexity scales linearly both in terms of the number of sequences and the number of epochs. With this refactoring in hand, we deliver a novel linear-time algorithm to evaluate the gradient of this (log-)likelihood wrt all epoch parameters simultaneously. Then we design and deploy an efficient HMC sampler that enables us to fit a large class of EBDS models in a Bayesian framework and provide an open-source implementation in the popular Bayesian Evolutionary Analysis by Sampling Trees (BEAST) software (Suchard et al. 2018).

Current approaches to Bayesian inference for EBDS epoch parameters employ a variety of prior assumptions to model the dependence structure between parameters across epochs. Some priors assume that birth, death and sampling rates across epochs are independent and identically distributed (iid) (Stadler et al. 2013, Gavryushkina et al. 2014, Vasylyeva et al. 2020). To smooth rate variation over time, temporally-auto-correlated priors such as Ornstein-Uhlenbeck smoothing prior (Du Plessis 2016), Gaussian Markov random fields (GMRF) priors (Condamine et al. 2018, Silvestro et al. 2019) and the horseshoe Markov random field for EBDS models (Magee et al. 2020) have been considered. Conveniently, both our linear-time gradients and our HMC approach generalize across all of these choices of prior without the need to construct model-specific sampling techniques and allow us to introduce the Bayesian bridge shrinkage prior to yield parsimonious time-varying rate patterns.

Across three real-world infectious disease examples that vary in the number of sequences, model dimension, and prior specification, we demonstrate the performance gain achieved by our implementation of an HMC transition kernel compared to random walk transition kernels. Moreover, for each of these datasets we infer key epidemiological parameters and demonstrate the utility of our scalable approach for providing reasonable estimates of pathogen transmission dynamics over time.

## 2 Methods

### 2.1 Setup

In an infectious disease setting, suppose an infected individual initiates an epidemic at time (measured backwards from the present day) *t*_*or*_ *>* 0, called the time of the origin. Then, each currently and newly infected individual disseminates the pathogen to others at a timevarying birth rate *λ*(*t*) and transitions into a noninfectious state at a time-varying death rate *μ*(*t*). At any given time, we may sample an infected individual with time-varying sampling rate *ψ*(*t*), at which point we add the time of sampling and a molecular sequence of their infectious agent into our time-stamped molecular sequence alignment **Y**. Further, we may posit *K* fixed time-points at which we randomly sample all infected individuals with associated vector of probabilities ***ρ*** = (*ρ*_1_, …, *ρ*_*K*_), adding the time and molecular sequence to **Y**. Note that this means that several individuals can be sampled at the same time point. The choice of the time-points is dependent on the dataset at hand and will be discussed later in this section. Every sampled infected individual may be treated and then become noninfectious with time-varying probability *r*(*t*) which we assume equal to one everywhere for complete sampling.

The model defined above provides a forward in time portrayal of the epidemiological process. Considering the *N* sampled and time-stamped sequences in **Y** as tree tips, there exists a (possibly unknown) phylogeny 𝒯 that depicts the evolutionary relationships among these sequences. Specifically, 𝒯 is a rooted, bifurcating tree with *N* tip nodes that correspond to the sampled sequences or their hosts from the population and *N* − 1 internal nodes that represent transmission events between hosts. We define the height of the nodes as the length of time between the time of the corresponding transmission/sampling events and the time of the most recent sampled sequence, which we refer to the present time, 0. Each node of 𝒯 is then associated with a node-height ≥ 0 relative to the present, such that the difference between the parent node-height and its child node-height is a branch length measured in the units of real time (e.g., years). We call the earliest internal node in 𝒯 the root and its node-height corresponds to the time of the most recent common ancestor (TMRCA). Therefore, we can further define the node heights of internal nodes to be bifurcation times and that of leave nodes to be sampling times. Accordingly, for a vector of bifurcation times, we have ***v*** = (*v*_1_, *v*_2_, …, *v*_*N*−1_) where *v*_1_ *< · · · < v*_*N*−1_. And we let ***u*** = (*u*_1_, *u*_2_, …, *u*_*N*_) be a vector of sampling times where *u*_1_ *< · · · < u*_*N*_.

For an episodic model, we make the assumption that all the rate parameters are piecewise constant across *K* different epochs with cut points ***t*** = (*t*_0_, …, *t*_*K*_), with *t*_0_ = 0 *< t*_1_ *< · · · < t*_*K*−1_ *< t*_*K*_. We also require *t*_*or*_ ≤ *t*_*K*_. Under this assumption, we can rewrite the time dependent birth rate *λ*(*t*) in terms of some unknown epoch-specific birth rate ***λ*** = (*λ*_1_, …, *λ*_*K*_), where *λ*(*t*) = *λ*_*k*_ for *t*_*k*−1_ *< t* ≤ *t*_*k*_. Similar parametrization applies to other parameters, so that we can express *μ*(*t*) in terms of ***μ*** = (*μ*_1_, …, *μ*_*K*_), *ψ*(*t*) in terms of ***ψ*** = (*ψ*_1_, …, *ψ*_*K*_) and *r*(*t*) in terms of ***r*** = (*r*_1_, …, *r*_*K*_). Without loss of generality, we let intensive sampling events happen at every time points in ***t***. Then we define ***ρ*** = (*ρ*_1_, …, *ρ*_*K*_), where *ρ*(*t*) = *ρ*_*k*_ for *t* = *t*_*k*−1_. We can remove these intensive sampling events at the epoch switching times from our model simply by setting ***ρ*** = **0**.

After reparametrizing the rates of the EBDS model, we can arrive at some key epidemiological quantities. For example, if we assume there are no intensive sampling events, we can specify the effective reproductive number as 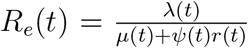. Other parameters that are important include the total rate of becoming noninfectious, which is defined as *δ*(*t*) = *μ*(*t*) + *ψ*(*t*)*r*(*t*), and the sampling proportion, defined as 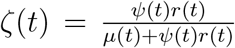. If we also assume removal of lineages upon sampling, these formulas can be further simplified by letting *r*(*t*) be constant and always equal to 1.

### 2.2 Probability Density of a Sampled Phylogeny

Recall we break time into intervals with cut points ***t*** = (*t*_0_, …, *t*_*K*_) defined by epochs. Within each epoch, we define a series of subintervals such that a new subinterval start at every bifurcation time ***v***, sampling time ***u*** and epoch switching time ***t***. We delineate the subinterval by indices *j*, which begins at *s*_*j*_ and terminates at *s*_*j*+1_ (where *s*_*j*_ *< s*_*j*+1_). If *t*_*or*_ = *t*_*K*_, then the grids ***s*** = (*s*_1_, …, *s*_2*N*−2+*K*_) can be obtained by joining the time points in ***v, u*** and ***t*** according to their ascending order when none of these times coincide with each other. If *t*_*or*_ *< t*_*K*_, we have *s*_2*N*−2+*K*_ = *t*_*or*_ instead of *t*_*K*_.

Consequently, each subinterval, inclusive on the left, is partitioned in such a way that it precludes the occurrence of an epoch switching, birth or sampling event within its boundaries. Within the *k*th epoch, the first subinterval starts at *s*_*j*_ = *t*_*k*−1_ and the last subinterval ends at 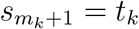. (Note for the last epoch *K*, the last subinterval ends at *t*_*or*_.) We assign *L*(*j*) to account for the number of lineages in 𝒯 that are extant in subinterval time (*s*_*j*_, *s*_*j*+1_].

Our likelihood derivation falls into the common framework with Stadler et al. (2013), Gavryushkina et al. (2014) and Magee & Höhna (2021). However, instead of writing the likelihood in terms of the times of node and epochs, we write it in terms of the subintervals *j*. This representation highlights the fact that the likelihood can be computed in one pass, starting at the present and ending at the origin. The interval-based representation of the likelihood is as follows:

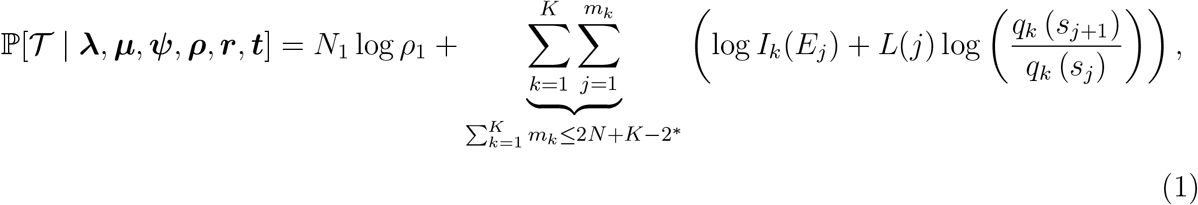

where *m*_*k*_ is the total number of subintervals in epoch *k*. (*: equality holds when no events happens at the exact same time except for the current).

The indicator function *I*_*k*_(*E*_*j*_) is labelled by the index *k*. This implies that the function is concerned with events occurring within the time frame (*t*_*k*−1_, *t*_*k*_]. We have *E*_*j*_ represent the event that takes place at the termination of subinterval *j* within epoch *k*. In most phylodynamic studies, ancestral sampling scenarios are not taken into account; therefore, our model is based on the assumption of a strictly bifurcating phylogenetic tree and does not involve considerations of ancestral sampling cases, which is distinctive from the work of Gavryushkina et al. (2014). Nonetheless, incorporating ancestral sampling into our framework is relatively straightforward. This can be achieved by setting the treatment probability to be less than 1 and adding the term *ψ*_*k*_(1 − *r*_*k*_) to our indicator function to account for events involving ancestral samples. Consequently, this indicator function takes the following form:

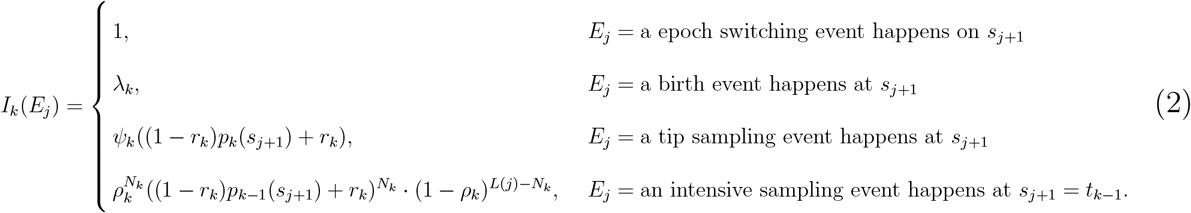

Note that *p*_*k*_(*t*) is the probability that an infected individual at time *t* has no sampled descendants when the process is stopped (i.e., at time *t*_0_), and *q*_*k*_(*t*) is the probability density of an individual at time *t* giving rise to an edge between *t* and *t*_*k*−1_ (not *t*_*k*_ since we define time to flow backwards which is the reverse of the generative process) for *t*_*k*−1_ *< t < t*_*k*_ in epoch *k*. We have *p*_0_(*t*_0_ = 0) = 1.

The intensive sampling probability at time *t*_*k*−1_ is *ρ*_*k*_ and the corresponding number of leaves sampled at that time is *N*_*k*_. The index here is intentionally misaligned to reconcile the fact that we model the epoch as left inclusive in time.

The definitions of the underlying functions, *q*_*k*_(*t*) and *p*_*k*_(*t*), follow the work from Stadler et al. (2013) and the detailed formulas are included in Supplementary Material S1. Note that our equation 1 does not condition the tree likelihood upon any particular properties, such as the presence of at least one sampled individual. Without loss of generality, additional conditioning schemes can be integrated by adding a factor to the log-likelihood; relevant discussions on this subject are available in Table S3 from the study by MacPherson et al. (2022).

As stated previously, our representation of the likelihood differs from the more standard nodewise representation (see for example Stadler et al. 2013, Gavryushkina et al. 2014, Wu 2014, Magee & Höhna 2021). Our representation makes it explicit that the likelihood computation can be accomplished in 𝒪(*N* +*K*) time (see Algorithm 1 for computational details). We demonstrate this behavior empirically in Supplementary Material S6. On the other hand, as we show in Supplementary Material S5, the conventional nodewise representation leads to ambiguities in the cost and a wide potentially range of computational complexities depending on implementation decisions. In Supplementary Material S6 we show empirically that formulations based on the nodewise representation include both implementations which are of the same computational order as ours (namely BEAST2 (Bouckaert et al. 2019) and RevBayes (Höhna et al. 2016)) and which scale worse in the number of epochs (TreePar (Stadler et al. 2013)).

### 2.3 Inference

In a Bayesan inference procedure, as introduced in Section 2.1, we use a multiple sequence alignment with the sampling times, the time-stamped sequences, **Y**, as the input data. Based on **Y**, we can form the posterior distribution over the product space of trees and EBDS model parameters as follows. First, a phylogeny 𝒯 is generated from the EBDS process defined in Section 2. Then we specify a molecular clock model that controls the rate at which evolution occurs on each branch of 𝒯. Under a molecular character-based CTMC substitution model, the columns in the sequence alignment evolve independently along the branches of the tree. Adoption of different substitution models is contingent upon the distinct attributes of the dataset under investigation (see Section 2.6.1). For the sake of notational convenience, we refer to the vector encompassing both substitution and clock model parameters as ***ω***. We denote by ℙ (***Y*** | ***ω***, 𝒯) the probability of the time-stamped sequences under the CTMC substitution model, known as the phylogenetic likelihood. Subsequently, we can factorize the posterior in the following manner:

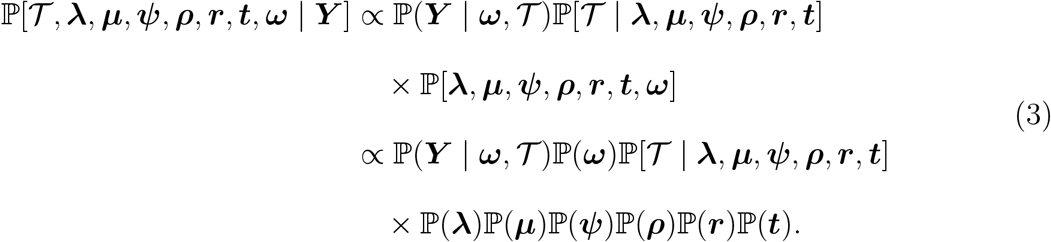

In phylodynamic analyses, it is sometimes advantageous to streamline the model by maintaining the death rate as constant. We can also presume the intensive sampling probability to be 0 and treatment probability to uniformly be 1 across all epochs. In handling time-varying parameters, we choose either iid priors or Markov random field models based on dataset-dependent assumptions pertaining to the patterns of change expected in rate parameters. In this paper, we specifically consider the GMRF and the Bayesian bridge Markov random field model, the latter of which we describe below.

With increasing complexity of the existing EBDS models, we seek to integrate Bayesian regularization methods to help manage the potentially vast quantity of model parameters. Specifically, we consider Markov random field priors which specify distributions on the incremental difference between the log-transformed rate parameters. By assigning a normal distribution to the incremental changes, we arrive at the GMRF priors that induce a smoothing effect on the change of rate parameters across contiguous epochs. This approach naturally leads to adjacent epochs exhibiting similar rate values. However, a strong data signal indicative of a rate change can still manifest in the resulting trajectory. By placing a heavy-tailed Bayesian bridge prior (Piironen & Vehtari 2017) on these, we achieve a more generalized extension of the GMRF model. The key distinction resides in the specification of the standard deviation arising from the normal priors on the increments. In this resulting Bayesian bridge Markov random field framework, each epoch’s increment is assigned an additional variable to account for variation, thereby affording greater flexibility to the model.

Supposing we have varied birth rates, we define the birth rate on the log scale 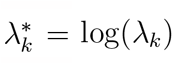. Then we have the prior on increments, 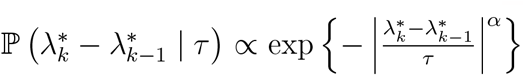 for *k >* 1, where *τ* is the global scale parameter that controls the overall degree of parameter variation. As *α* diminishes, the function 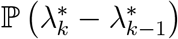 accrues an increased density close to zero. For the purpose of our study, we establish *α* = 0.25 to address a potent prior assumption that 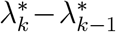 is proximate to 0 without inducing any problems related to mixing issues. In other words, we do not anticipate substantial fluctuations in the birth rates across consecutive epochs (but allow for rapid rate shift, for example during the exponential growth phase.) Another important parameter is the local scale, denoted as *ν*_*k*_, which is specific to an individual increment 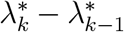. Its density regulates the magnitude of the spike and the tail behavior of the above marginal 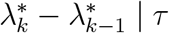.

Note that the GMRF model can be perceived as a specific instance of the Bayesian bridge MRF, where all the local scale parameters are equalized to 1 and *α* is fixed at 2. In this case, the increment differences adhere to a normal distribution whose variance is solely governed by a single global scale parameter.

To complete our model, a normal prior is assigned to 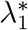 in adherence with the method outlined in Magee et al. (2020). We obtain the mean parameter of the prior using an empirical Bayes method. This provides a crude estimate of the log rate parameter, coupled with a standard deviation that is sufficiently large to encompass all possible values (See S3). We apply a Gamma(1,1) prior to *ϕ* = *τ*^−*α*^. This selection is grounded on a combination of theoretical considerations and empirical validation and allows for an efficient Gibbs sampler for *τ*.

To regularize the tail behavior, we leverage the shrunken-shoulder version of the Bayesian bridge prior and limit the bridge to have light tails past the slab width, *ξ* (Piironen & Vehtari 2017, Nishimura & Suchard 2023). An efficient update of Markov random field models global and local scale parameters (for Bayesian bridge priors) follows Nishimura & Suchard (2023). In this framework, the prior on the increment space represented as a scale mixture of normal distributions:

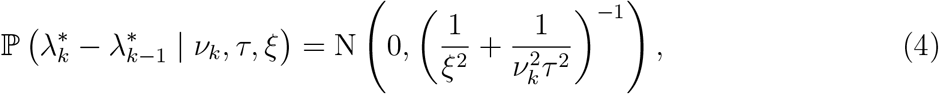

where *ν*_*k*_ is called the local scale parameter and *τ* is the global scale parameter. (Note that *ν*_*k*_ has an exponentially tilted stable distribution with characteristic exponent *α/*2.) This mixture representation aids in clarifying the local adaptivity of the Bayesian bridge prior as considerable changes in rates can be accommodated by an increase in *ν*_*k*_ without necessitating a rise in *τ*. The inclusion of the slab width helps to bound the variance of increments to *ξ*^2^. We set *ξ* = 2, which creates a reasonable upper limit on the variations in birth rate between consecutive epochs.

In our study, we primarily focus on sampling ℙ [𝒯 | ***λ, μ, ψ, t***]. With increasing numbers of epochs, the parameter space of the EBDS model expands quickly, exhibiting substantial correlation between adjacent epochs. To improve the sampling efficiency, we utilize HMC method to concurrently sample the time varying model parameters and ensure a high acceptance rate.

### 2.4 Hamiltonian Monte Carlo Sampling

Hamiltonian Monte Carlo is a widely-used Markov chain Monte Carlo method to sample from a target distribution effectively. Given a target parameter ***θ*** with a posterior probability density *π*(***θ***), HMC iteratively generates samples from the target distribution by simulating the dynamics of a physical system whose equilibrium distribution is equal to *π*(***θ***). In particular, HMC introduces an auxiliary momentum parameter ***d***, which is typically chosen to follow a multivariate normal distribution with zero mean and covariance matrix ***M***, i.e., ***d*** ∼ *N* (0, ***M***). ***M*** is also known as the mass matrix, which serves as a hyperparameter. The Hamiltonian function of the system is defined as:

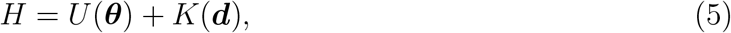

where *U* (***θ***) = − log(*π*(***θ***)) is the potential energy, and *K*(***d***) = ***d***^T^***Md*** is the kinetic energy of the system.

Starting from the current state (***θ***_0_, ***d***_0_), HMC updates the state according to the following differential equations:

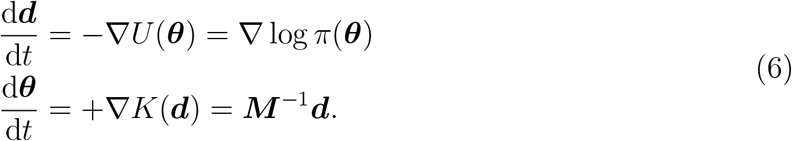

The simple and effective “leapfrog” method (Neal et al. 2011) approximates the solution to (6) numerically:

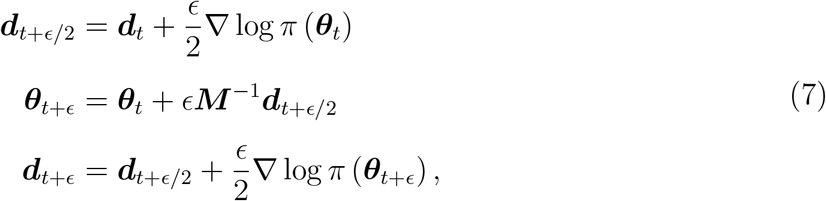

where *ϵ* is the size of each leapfrog step, and *n* steps are required to simulate the Hamiltonian dynamics from time *t* = 0 to *t* = *nϵ*. In practice, the “leapfrog” method has been shown to be stable and accurate for a wide range of step sizes (Neal et al. 2011).

The default choice of the mass matrix is the identity matrix. However, using a different ***M***, such as a log-posterior Hessian approximation can largely enhance the efficiency of HMC sampling. In this work, ***M*** is adaptively tuned to estimate the expected (diagongal) Hessian averaged over the prior distribution. This design choice alleviates some computational burden, following the work of Ji et al. (2020).

### 2.5 Gradient

HMC sampling of the model parameters requires the gradient of the log-likelihood derived from (1) wrt the EBDS model rate parameters. The gradient is the collection of derivatives wrt model parameters:

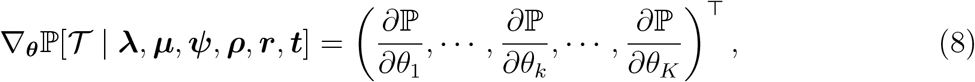

where *θ*_*k*_ *∈ {λ*_*k*_, *ψ*_*k*_, *μ*_*k*_, *ρ*_*k*_*}* is a unified parameter to reduce notation clutter.

Given the piece-wise constant nature of the model, the likelihood assumes a consistent form across all epochs. Therefore, we can examine the gradient of the log-likelihood at each epoch separately. We denote the log-likelihood at epoch *k* and phylogeny segment *j* as:

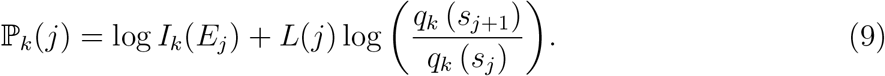

We can further get individual terms in (8) by accumulating contributions from each epoch and the corresponding phylogeny segments:

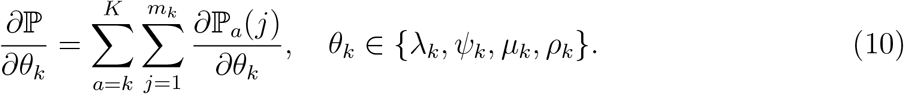

By examining the interdependency between epochs, we discern that a given epoch *k* exerts influence on the gradient of parameters pertaining to that and all preceding epochs. Consequently, it becomes necessary to consider 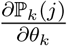 and 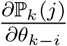 respectively, where *i* is a positive integer ranging between 1 and (*k* − 1).

First, we consider the gradient contribution at epoch *k* wrt the current model parameters 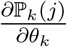, where *θ*_*k*_ *∈ {λ*_*k*_, *ψ*_*k*_, *μ*_*k*_, *ρ*_*k*_*}*.

Then we have the following cases:

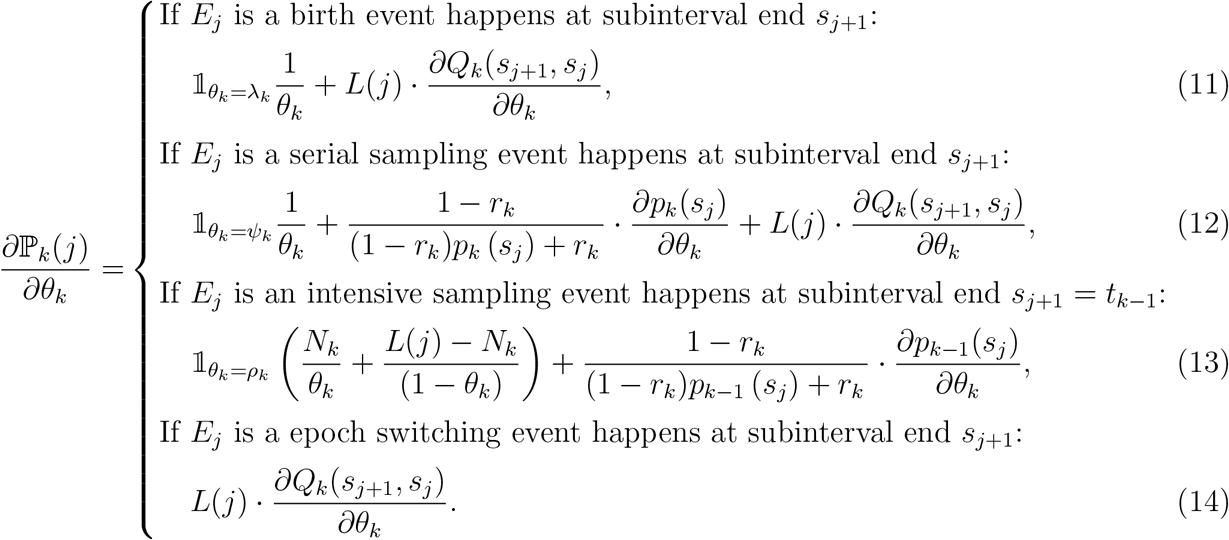

Note that 𝟙 is the indicator function. We leave the explicit expression of the shared terms in (11)-(14) to Supplementary Material S2.

Second, we consider the gradient at epoch *k* wrt the previous model parameters 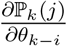, where *θ*_*k*−*i*_ *∈ {λ*_*k*−*i*_, *ψ*_*k*−*i*_, *μ*_*k*−*i*_, *ρ*_*k*−*i*_*}*:

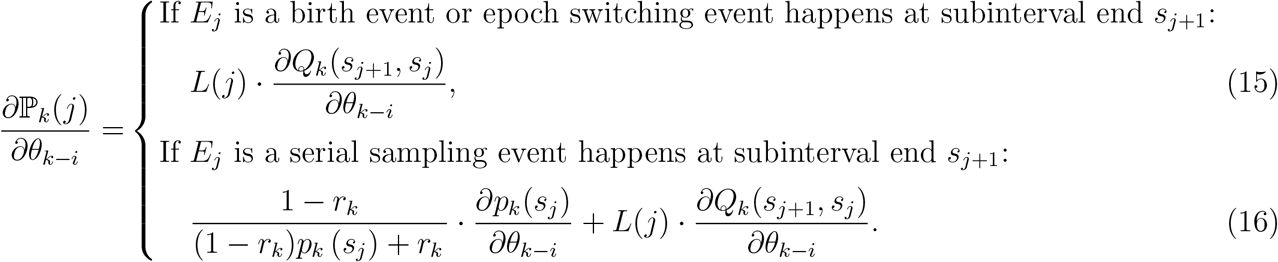

We also leave the explicit expression of the shared terms in (15)-(16) in Section S2.

Third, we discuss the gradient at epoch *k* wrt the treatment probability **r**. In (1), the treatment probabilities at different epochs only affect the current epoch. Therefore, we only need to consider 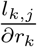 as follows:

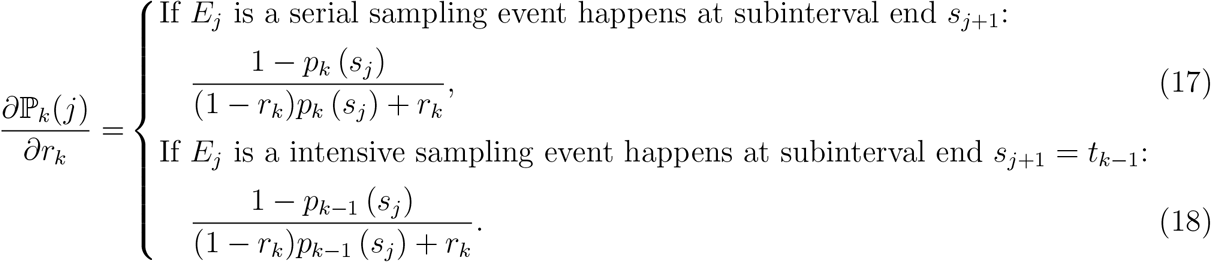

The total gradient wrt **r** can be obtained similar to (10).

To determine the computation complexity of gradient evaluation, we can assume the gradient calculation for 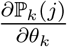 takes constant time. The model has *K* epochs, where each epoch has 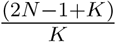 phylogeny segments in average. According to (10), the total computation complexity is 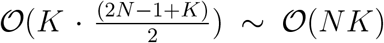, since *K* ≪ *N*. We demonstrate this result through a series of timing experiments presented in Supplementary Material S6 where we also compare the efficiency of gradients calculations with the automatic differentiation algorithm implemented in the VBSKY (Ki & Terhorst 2022) package based on JAX library (Bradbury et al. 2018). Figure S5 shows our analytical gradients implemented in BEAST significantly outpace the VBSKY method.

### 2.6 Analysis

#### 2.6.1 Examples

We evaluate the relative effectiveness of MH-MCMC and HMC transition kernels under the EBDS model using three phylodynamic examples. The first example comprises 274 sequences of the Pol locus of HIV-1 subtype A sampled in Odesa, Ukraine from 2000 to 2020 that Vasylyeva et al. (2020) previously analyzed to assess the population-level impact of the transmission reduction intervention project (TRIP) on HIV transmission (Nikolopoulos et al. 2016). Following this previous analysis, we establish a cutoff point of 50 years for the EBDS model. Within this period of time, we let the birth, death and sampling rates vary across 10 epochs mirroring the grid points specified by Vasylyeva et al. (2020). Note that for better comparability to the original work (Vasylyeva et al. 2020), we place iid lognormal priors on the rate parameters. Both the previous and our analysis assume an HKY nucleotide substitution (Hasegawa et al. 1985) model with discrete-gamma-distributed rate variation among sites (HKY+G) (Yang 1994), and an uncorrelated lognormal relaxed molecular clock model (Drummond et al. 2006) (UCLD), with a CTMC rate-reference prior (Ferreira & Suchard 2008) on the clock-model mean, truncated between 1 *×* 10^−3^ 3 *×* 10^−3^, and a normal prior (with mean = 5 *×* 10^−4^ and standard deviation = 5 *×* 10^−4^) on the standard deviation. We use a normal distribution prior (with mean = 35, standard deviation = 5) on the time to the most recent common ancestor, in accordance with the previous study.

Second, we examine the transmission dynamics of 637 human influenza A/H3N2 HA genes across 12 epidemic seasons sampled from New York state Rambaut et al. (2008) following the study of Parag et al. (2020). We set an EBDS model cutoff value of 13 years and infer time-varying birth and sampling rates across 78 epochs, each representing 2 months in time, and a constant-over-time death rate. Preceding studies focused on the evolutionary dynamics of influenza A/H1N1 virus mostly utilize the coalescent models. These studies predominantly rely on Gaussian process smoothing (Karcher et al. 2020, Bhattacharjee et al. 2023). Following the same path, we seek to use GMRF prior distributions for the birth and sampling rates. Our approach accommodates the considerable variability in the effective reproductive number across different flu seasons from 1993 to 2005. We adopt the same substitution and clock models from Rambaut et al. (2008). Specifically, to account for potential differences in the rate of substitution between the first and second codon positions compared to the third, we employ the SRD06 substitution model (Shapiro et al. 2006) and apply an HKY nucleotide substitution model with discrete-gamma distributed rate heterogeneity for both codon-position partitions (1st + 2nd, and 3rd). We further assume a UCLD clock model and employ the default priors from BEAST on the substitution and clock model parameters.

Lastly, to demonstrate the potential our linear-time algorithms afford phylodynamic analyses on larger data sets, we examine 1610 full Ebola virus (EBOV) genomes sampled between 17 March 2014 and 24 October 2015 from West Africa (Dudas et al. 2017) to explored the factors contributing to the spread of Ebola during the 2014-2016 epidemic. We set a EBDS model cutoff value of 2 years and infer time-varying birth and sampling rates for 24 epochs, each corresponding to a month in time, and a constant death rate. For choosing the priors on the rate parameters, we incorporate information from previous studies on the transmission dynamics of Ebola virus disease in West Africa (Fang et al. 2016, Nyenswah et al. 2016). The number of confirmed cases first persisted at a relatively low level and started to soar in the mid-Summer of 2014, followed by a consistent peak and a dramatic decrease after the initiation of some key intervention events. Considering the potential fast shifts projected to the effective reproductive number, we apply the Bayesian bridge MRF model as the prior for the incremental differences in the birth and sampling rates. Based on Dudas et al. (2017), we assume a HKY+G substitution model independently across four partitions (codon positions 1, 2, 3 and non-coding intergenic regions) and a log-normally-distributed relaxed molecular clock model with a CTMC reference prior on the clock model mean, and leave all other priors on substitution and clock model parameters at their BEAST defaults.

#### 2.6.2 Implementation

We conduct all analyses using extensions to BEAST 1.10 (Suchard et al. 2018) and the highperformance BEAGLE 4.0 library (Ayres et al. 2019) for efficient computation on central processing units (CPUs). We take the timing measurements using a Macbook Pro equipped with an M1 Pro chip that features 8 CPU cores and 32GB of RAM. For all experiments involving BEAST, we utilized the Azul Zulu Builds of OpenJDK version 18 on the ARM architecture.

To compare the performance of the two transition kernels in estimating the EBDS model parameters, we conduct efficiency comparison analyses that focused solely on the estimation of the birth-death model’s rate parameters. Specifically, we fix the phylogeny to the maximum clade credibility (MCC) tree, a tree with the maximum product of the posterior clade probabilities summarized from the Bayesian joint phylogeny inference. We analyze all data sets using BEAST with logging performed every 1000 iterations. We run our algorithm on the HIV example for 300 million iterations when using MH-MCMC transition kernel and 30 million iterations for HMC transition kernel. Also, to obtain convergent results for the influenza example, we run analyses using MH-MCMC and HMC transition kernels for 300 million and 50 million states, respectively. For the Ebola example, we run analyses using MH-MCMC and HMC transition kernels for 100 million and 30 million states, respectively. For all analyses, we discard 10% of the MCMC chain samples as burn-in.

We calculate the effective sample size (ESS) for each rate parameter of interest using the coda package (Plummer et al. 2006) in CRAN (R Core Team 2021). ESS quantifies the degree to which auto-correlation within MCMC iterations contributes to uncertainty in estimates (Ripley 2009). We average ESS per compute-hour for each parameter across 10 independent runs to reduce Monte Carlo error in each estimate, aiming for a maximal Monte Carlo error of 10%. We report the relative increase in ESS per hour of the HMC sampler compared with the MH-MCMC sampler over all rate parameters.

We also conduct phylodynamic analysis for each of the three examples under a joint phylogeny inference scheme to mitigate potential bias from the fixed phylogeny, following the model specifications discussed in Section 2.6.1. Under these settings, we simulate MCMC chains for all examples of’ 500 million iterations using HMC transition kernel with logging performed every 1000 iterations.

## 3 Results

### 3.1 Performance Improvements

Figure 2 shows the binned ESS per hour estimates of the EBDS model rates (*λ, μ, ψ*) that the MH-MCMC and HMC samples generate for all three viral examples. Table 1 summarizes the performance improvements by reporting the relative increase in the minimum ESS per hour comparing both samplers across all rate parameters.

**Table 1:** Relative speedup in terms of effective sample size per hour (ESS/h) of HMC Over MH-MCMC for all three data Sets from fixed phylogeny analyses.

| Example | Minimum ESS/h |  | HMC<br>Speedup |
| --- | --- | --- | --- |
|  | MH-MCMC | HMC |  |
| HIV (10 epochs) | $1.60 \times 10^1$ | $3.91 \times 10^3$ | $2.45 \times 10^2$ times |
| Influenza (78 epochs) | $2.43 \times 10^{-1}$ | $1.93 \times 10^1$ | $7.94 \times 10^1$ times |
| Ebola (24 epochs) | $5.30 \times 10^0$ | $6.73 \times 10^1$ | $1.27 \times 10^1$ times |

**Figure 1:**
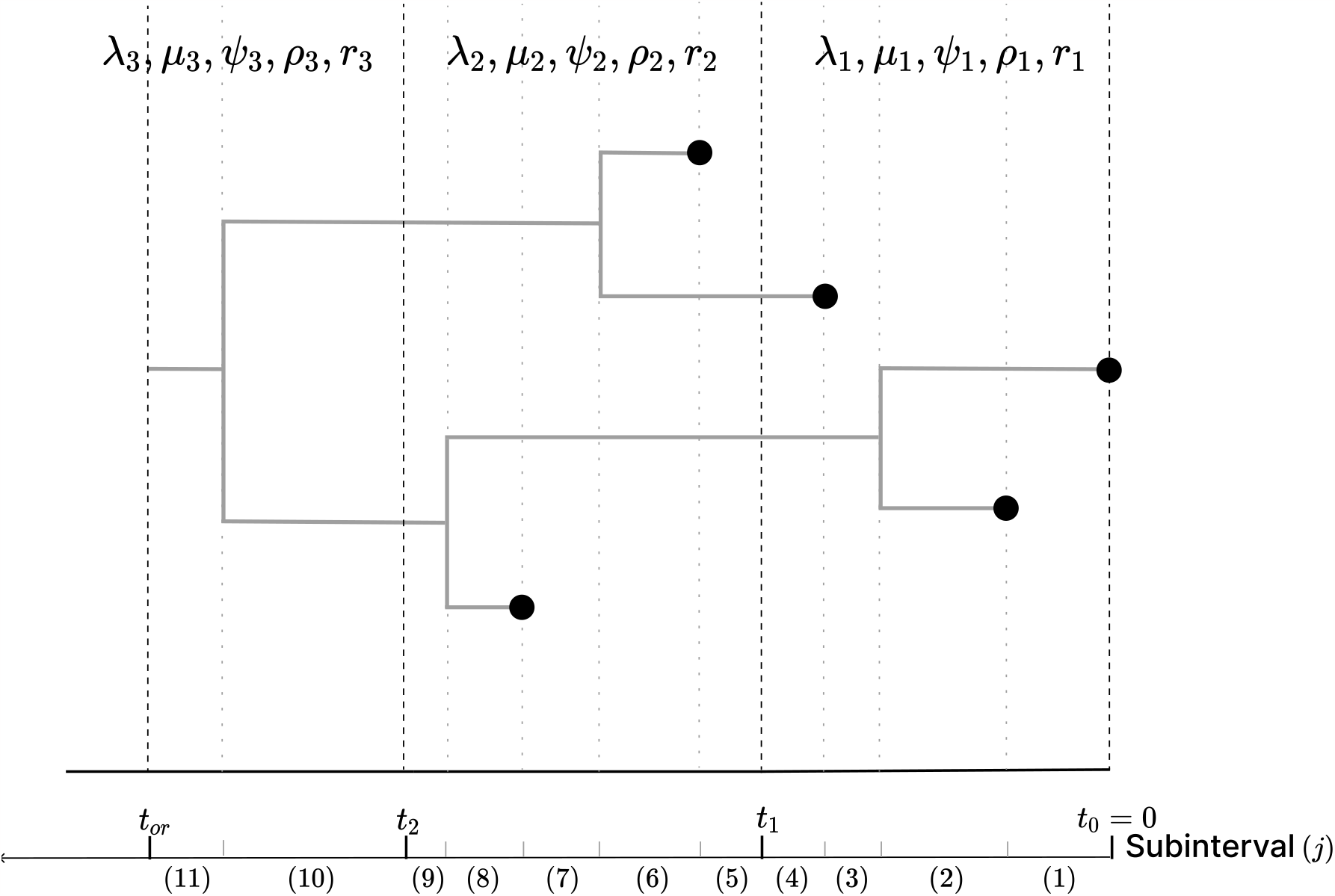
A phylogeny arising from an EBDS model. This sampled phylogeny has three epochs (with epoch switching time *t*_1_, *t*_2_) and thus three sets of model parameters including rates and probabilities. For every epoch, each branch is further divided into subinterval that starts at *s*_*j*_ and ends at time *s*_*j*+1_ so that no epoch switching, birth or sampling event occurs within it. Each subinterval within each epoch *k* is represented by a phylogeny segment index, *j*.

**Figure 2:**
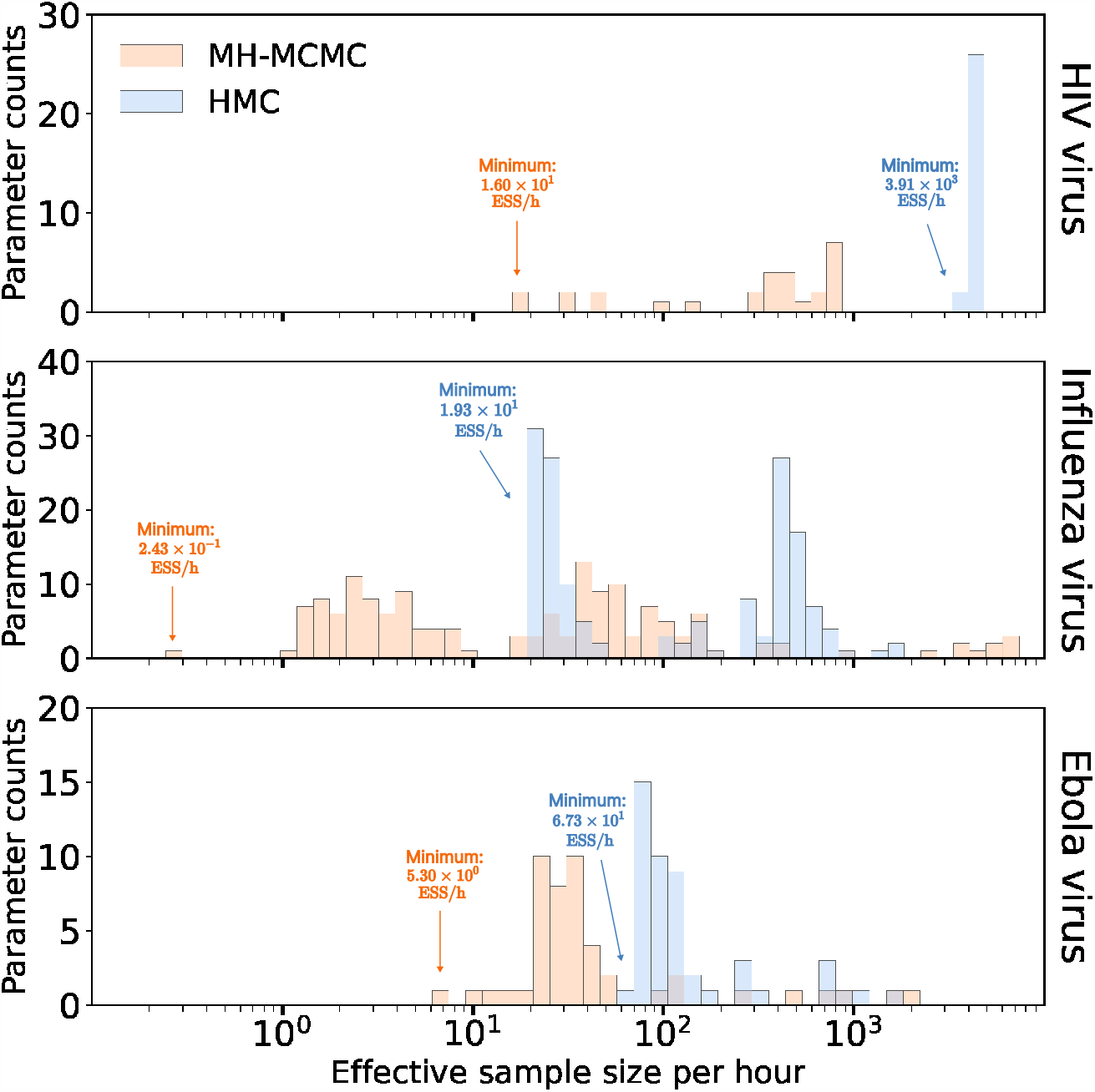
Efficiency Comparison between random walk Metropolis-Hastings (MH-MCMC) and Hamiltonian Monte Carlo (HMC) samplers. Bars correspond to the estimated effective sample size per hour averaged across 10 independent runs for all rate parameters. The height of each bar indicates the number of parameters that achieve the given ESS per hour value.

The HIV example assumes that time-varying rates are *a priori* independent across epochs and HMC demonstrates an approximate 245-fold acceleration relative to MH-MCMC. Likewise, the influenza example imposes a GMRF across epochs and returns an approximate 79.4-fold speed-up. On the other hand, the EBOV example enforces heavier shrinkage, and hence higher *a priori* correlation between epochs, and yields a smaller yet computationally impactful (approximately 12.7-fold) performance increase.

### 3.2 HIV dynamics in Odesa, Ukraine

In the context of conducting phylodynamic analyses using EBDS models, we are primarily interested in the value and trend of effective reproductive number over time *R*_*e*_(*t*) that is the average number of secondary cases per infectious case in a population made up of both susceptible and non-susceptible hosts. If *R*_*e*_ *>* 1, the number of cases is growing, such as at the start of an epidemic; if *R*_*e*_ = 1, the disease is endemic; and if *R*_*e*_ *<* 1, there is an expected decrease in transmission (Nishiura & Chowell 2009). Under the EBDS model, given the absence of intensive sampling events, if an individual becomes infected at time *t*, we can use the rate parameters at time *t* to obtain an estimated 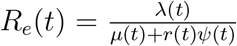. Furthermore, in all our analyses for infectious disease phylodynamics, we maintain *r*(*t*) = 1 as constant. This assertion carries the assumption that upon diagnosis and sequencing, an individual ceases to be a source of infection. This could be due to treatment, death, or geographical relocation, rendering them incapable of onward transmission.

To assess the effects of TRIP for reducing the transmission of HIV in Odesa, we fit the EBDS model with varying birth, death and sampling rates and plot the resulting *R*_*e*_(*t*) trend estimate in Figure 3. We apply iid lognormal priors on the rate parameters to stay consistent with the methods in previous study (Vasylyeva et al. 2020).

**Figure 3:**
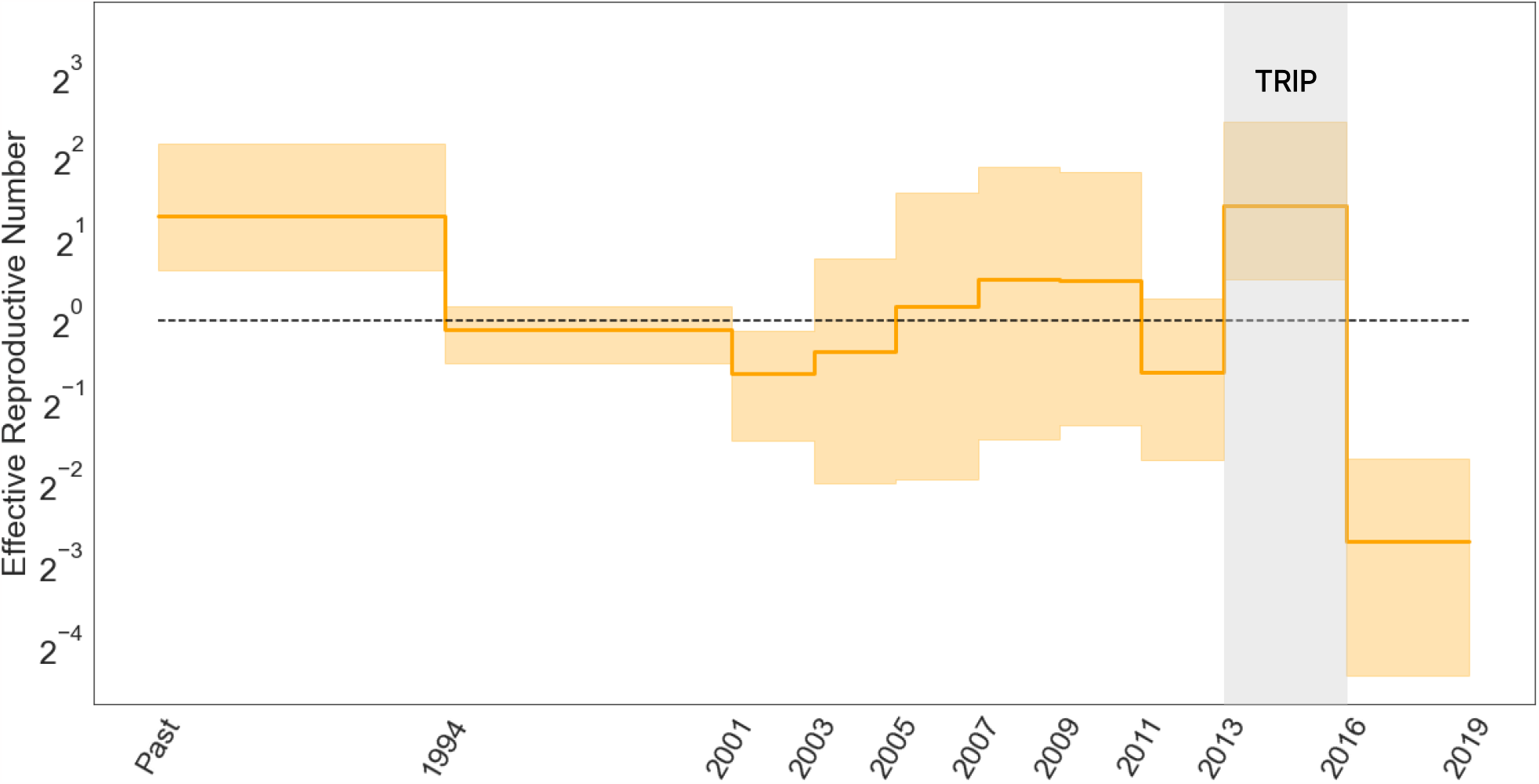
Posterior median (solid line) and 95% credible intervals (CI) indicated by the shaded areas of the effective reproductive number estimates (*R*_*e*_) through time for HIV epidemic in Odesa, where the black dotted line represents the epidemiological threshold of *R*_*e*_(*t*) = 1.

Estimates of *R*_*e*_(*t*) appear mostly to accord with previous findings that identify a drop in infection rate subsequent to the implementation of the TRIP intervention. Focusing on the period from 2013 to early 2016, when TRIP was enacted, our posterior mean estimate of *R*_*e*_ is 2.64 (95 % CI: 1.18 - 5.43); while post-intervention, the posterior mean reduces to 0.152 (95 % CI: 0.03 - 0.32). This latter value, falling below the critical threshold of 1, signifies the potential deceleration of HIV transmission.

### 3.3 Seasonal Influenza in New York State

While influenza viruses circulate throughout the year, peak influenza outbreaks in the United States typically occurs between December and February. Rambaut et al. (2008) employed a non-parametric coalescent model to elucidate the cyclical patterns of variation in the population size, uncovering a notable increase in genetic diversity at the beginning of each winter flu season. Subsequently, Parag et al. (2020) demonstrated that incorporating sampling intensity into the otherwise sampling-naive non-parametric coalescent process improves the precision of these inferred cycles. With a GMRF smoothing prior on increments, our model also offers the potential for accurately inferring seasonal behaviour and achieving the precision of parameter estimations.

Figure 4 presents posterior estimates of the effective reproductive number *R*_*e*_(*t*) for the alignment of 637 A/H3N2 HA sequences from New York state. As expected, the trajectory is highly cyclic, and all peaks lie near the midpoint of the influenza seasons with estimated *R*_*e*_ larger than 1. For the 2000/2001 and 2002/2003 seasons, where almost all infections were attributed to other sub-types of influenza viruses as indicated by the surveillance data and previous work (Centers for Disease Control and Prevention n.d., Parag et al. 2020), we observe the 95% CI of the estimated peak cover values from 0.68 to 1.3 and from 0.48 to 1.4, respectively. This suggests that their true *R*_*e*_ values might have fallen below 1. Similar to the results given by the non-parametric coalescent with sampling analysis (Parag et al. 2020), we capture a minor peak in the 1995/1996 season, where the inferred *R*_*e*_ is slightly above one. This again echoes with the fact that the influenza case composition during the 1995/1996 season was characterized by a mix of A/H1N1 and A/H3N2 infections (Ferguson et al. 2003). This diversity in infection types led to a less significant elevation in the effective reproductive number for that specific year.

**Figure 4:**
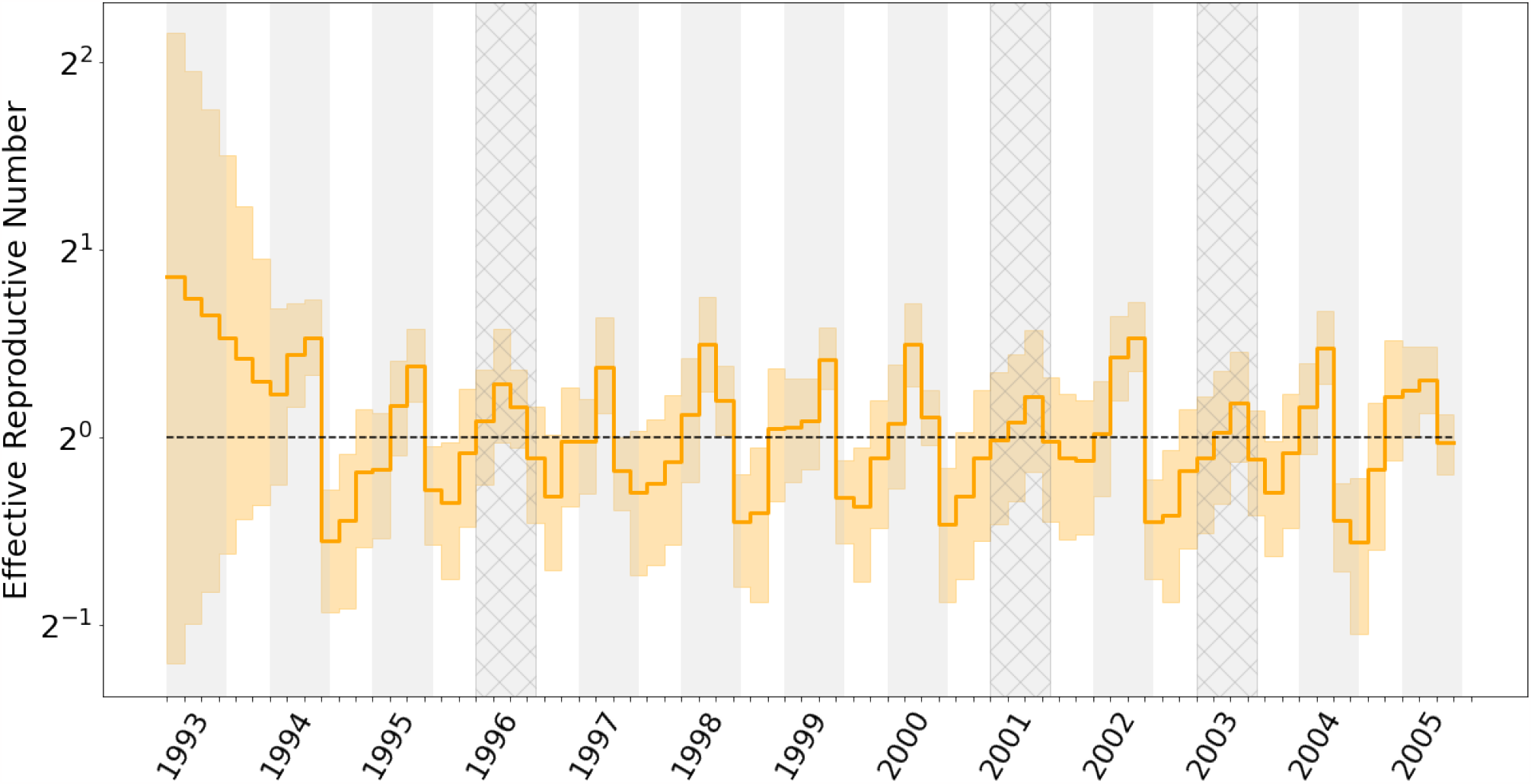
Median (solid orange line) and 95% credible intervals indicated by the shaded orange areas for the effective reproductive number estimates (Re) through time. Gray shading in the graph represents the rough duration of influenza monitored in New York state for each season, spanning from epidemiological week 40 to week 20 of the following year. Seasons where A/H3N2 was not the dominant influenza virus subtype are cross-hatched.

**Figure 5:**
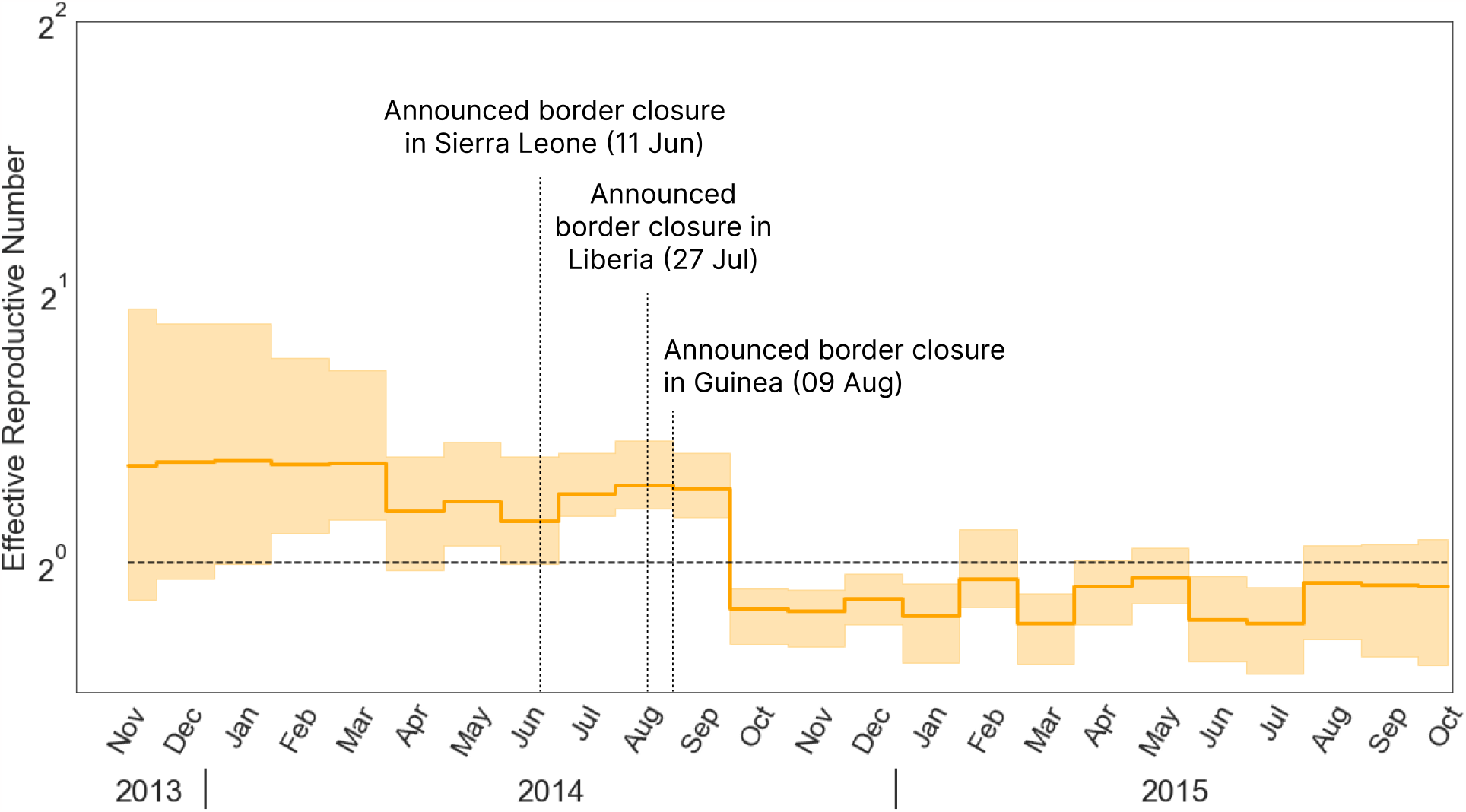
Median (solid line) and 95% credible intervals indicated by the shaded areas of the effective reproductive number estimates (*R*_*e*_) through time for Ebola outbreak in west Africa. The black dotted line represents the epidemiological threshold of *R*_*e*_ = 1.

### 3.4 Ebola epidemic in West Africa

Using EBDS model assisted by the HMC sampler, we are able to analyze the 2014 Ebola epidemic in West Africa using the full 1610-sequence alignment and metadata of sampling times taken from the work by Dudas et al. (2017). Previously, researchers have applied birthdeath models extensively for the phylodynamic analysis of the Ebola outbreak. Stadler et al. (2014) adopted a series of birth death models to capture the early trend of the infection of Ebola virus in Sierra-Leone. They used 72 Ebola samples from late May to mid June 2014 with three epochs, and estimated the corresponding effective reproductive number in each period. Zhukova et al. (2022) applied the multi-type birth death models to the 1610 sequence data. However, their analysis was based on the maximum likelihood estimation. To demonstrate the scalability of our method, we also take the 1610 sequence data and fit the EBDS model with 24 epochs for a finer time resolution to provide more precise estimation of the effective reproductive number. Here, we employ a Bayesian bridge MRF prior on rate increments to avoid spurious rate variations while capturing significant rate shifts.

Our inference results give an estimated posterior mean effective reproductive number at the beginning of the epidemic before December 2013 as 1.65 (95 % CI: 0.41 - 3.05). Dudas et al. (2017) show that after the international border closure of Sierra Leone on 11 June 2014, followed by Liberia on 27 July 2014, and Guinea on 9 August 2014, the relative contribution of international border to overall viral migration is significantly lower. The change-point probability is the highest from August to September. This finding stands clearly compatible with our EBDS inference that demonstrates a drop of *R*_*e*_ from 1.3 (posterior mean, 95 % CI: 1.01 - 1.59) to 0.79 (95 % CI: 0.62 - 0.91) after September 2014 when the international travel restrictions are in place across the three countries.

## 4 Discussion

Birth-death models serve as fundamental tools for modeling the temporal progression of epidemics. In extending the work of Stadler et al. (2013), Gavryushkina et al. (2014), we have provided a systematic representation of the EBDS model for phylodynamics that promotes scalability. Our general re-formalization of the EBDS likelihood identifies that its computation is simply 𝒪 (*N* + *K*), foreshadowing an 𝒪 (*NK*) algorithm to deliver its gradient wrt time-varying birth, death or sampling rates across *K* epochs. This optimal scaling enables HMC sampling to more efficiently explore the high-dimensional joint distribution of rates as we increase the number of sequences and the number of model epochs to learn these processes at a finer time-resolution. HMC also emits an agnostic approach to incorporate a variety of prior assumptions about these time-varying trends, without the need to hand-craft specialized transitions kernels for specific priors. Moreover, as suggested by Ji et al. (2020), we take measures to enhance the efficiency of our HMC sampler by preconditioning the mass matrix based on the Hessian of the log-prior.

Through three viral epidemic examples, we show that our HMC-assisted approach considerably accelerates Bayesian inference across three very different choices of prior models. Our preconditioned HMC sampler achieves roughly 10- to 200-fold increase over the widely used MH-MCMC sampler in terms of the minimum ESS per unit-time. The enhanced efficiency gains are particularly beneficial given the increasing use of phylodynamic inference techniques in conducting real-time evaluations of outbreak patterns.

For applying our model in phylodynamic analyses of disease epidemics, we first examine our EBDS model on the effects of TRIP for reducing the transmission of HIV in Ukraine, and our inference results support a decreased rate of transmission following the TRIP intervention. Applied to seasonal Influenza in New York city, our model is able to accurately capture the complex pattern of variation in *R*_*e*_ during each influenza season. Applied to the Ebola outbreak in West Africa, our model supports the effect of international travel restrictions characterized as a noticeable decrease in *R*_*e*_ following the border closure of the three countries in West Africa.

In the EBDS model, Stadler and colleagues (Stadler et al. 2013) have indicated that the three rate parameters, *λ, μ*, and *ψ*, cannot be simultaneously identified. This issue of unidentifiability in complex birth-death processes has also been recently discussed by Louca & Pennell (2020). In our own empirical analysis, problems related to unidentifiability seldom manifest when we restrict ourselves to estimating no more than two time-varying rate parameters. Instead, the primary challenge appears to be the multimodal nature of the posterior distribution. Legried & Terhorst (2022) have demonstrated that, under certain conditions, piecewise constant birth-death models can be reliably inferred and differentiated. Furthermore, Kopperud et al. (2023) showed that rapidly changing speciation or extinction rates can be accurately estimated. This lends credence to the identifiability of patterns we observed in our phylodynamic analysis of pandemics such as the seasonal influenza and the Ebola outbreaks.

Current methods to estimate the expected Hessian averaged over the posterior distribution improves upon the previous work (Girolami & Calderhead 2011) by avoiding excessive computational burden. However, it relies on numerical approximations to compute the Hessian, leaving room for potential performance enhancements. To further optimize the methodology, we can advance beyond analytical solutions solely for gradients and extend them to encompass the analytical Hessian. This would smooth the path of updating the adaptive mass matrix, offering opportunities for better outcomes in terms of both efficiency and accuracy.

In many scenarios, the examination of EBDS models is contingent upon having some preliminary understanding of how to identify the epoch switching time and the length of duration of each epoch. However, it is possible that information available through epidemiological surveillance is insufficient. Moreover, the choice of epoch duration can be related to the uncertainty in the timing of the rate shifts (Magee et al. 2020). In this study, our strategy aims to increase the number of epochs and leverage regularizing priors, striving to achieve a refined grid of timelines. Nevertheless, constraints persist on the maximum epochs feasible with our HMC algorithm, particularly when confronted with computational limitations or models exhibiting multimodality challenges. One possible solution entails simultaneously inferring epoch duration, epoch switching times, and rate parameters via the reversible-jump MCMC method (Wu 2014). However, this method requires one to integrate across models with differing dimension, which demands substantial effort and might be impractical for large datasets.

Considering these cases, if the piece-wise constant model assumptions can be lifted so that we can obtain a smoothly differentiable likelihood function, it would inherently aid in deriving gradients concerning node ages and epoch switching times. This advancement would, in turn, improve our current implementation, empowering us to infer, rather than presuppose, epoch switching times, with enhanced scalability prospects. It would also enhance the sampling efficiency from joint phylogeny posterior distributions, by enabling us to take advantage of recent work by Ji et al. (2021), yielding a pronounced improvement in the analytical capacity of our models.

In anticipation of future advancements that will improve upon standard HMC methods and broaden the applicability of the current EBDS model, we present a comprehensive framework in this manuscript. This framework facilitates phylodynamic analysis on largescale sequence data and employs regularization techniques to yield a finely-resolved, regular grid that effectively aids in our understanding of the impact of the pandemics.

## 5 Acknowledgements

This work was supported through National Institutes of Health grants U19 AI135995, R01 AI153044 and R01 AI162611. We gratefully acknowledge support from Advanced Micro Devices, Inc. with the donation of parallel computing resources used for this research.

## Supplementary Material

### S1 Likelihood Derivation

#### S1.1 Formulas for Likelihood Related Functions

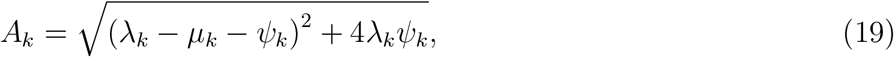

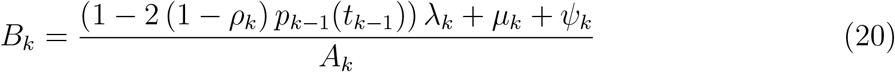

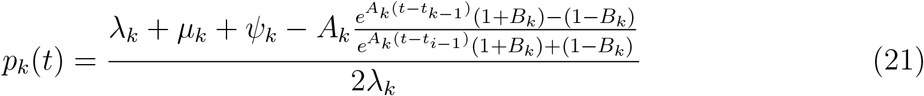

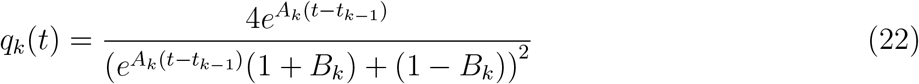

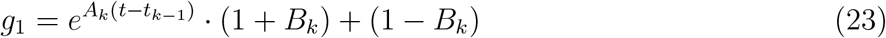

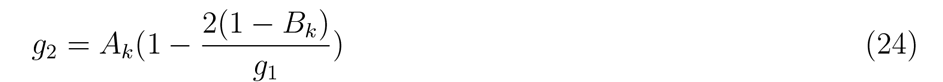

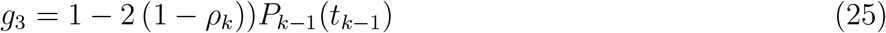

#### S1.2 Implementation Algorithm

Detailed algorithm for likelihood calculation is shown below based on the equations listed in Section 2.2 of the main text and from the section above.

##### Algorithm 1

Likelihood Calculation

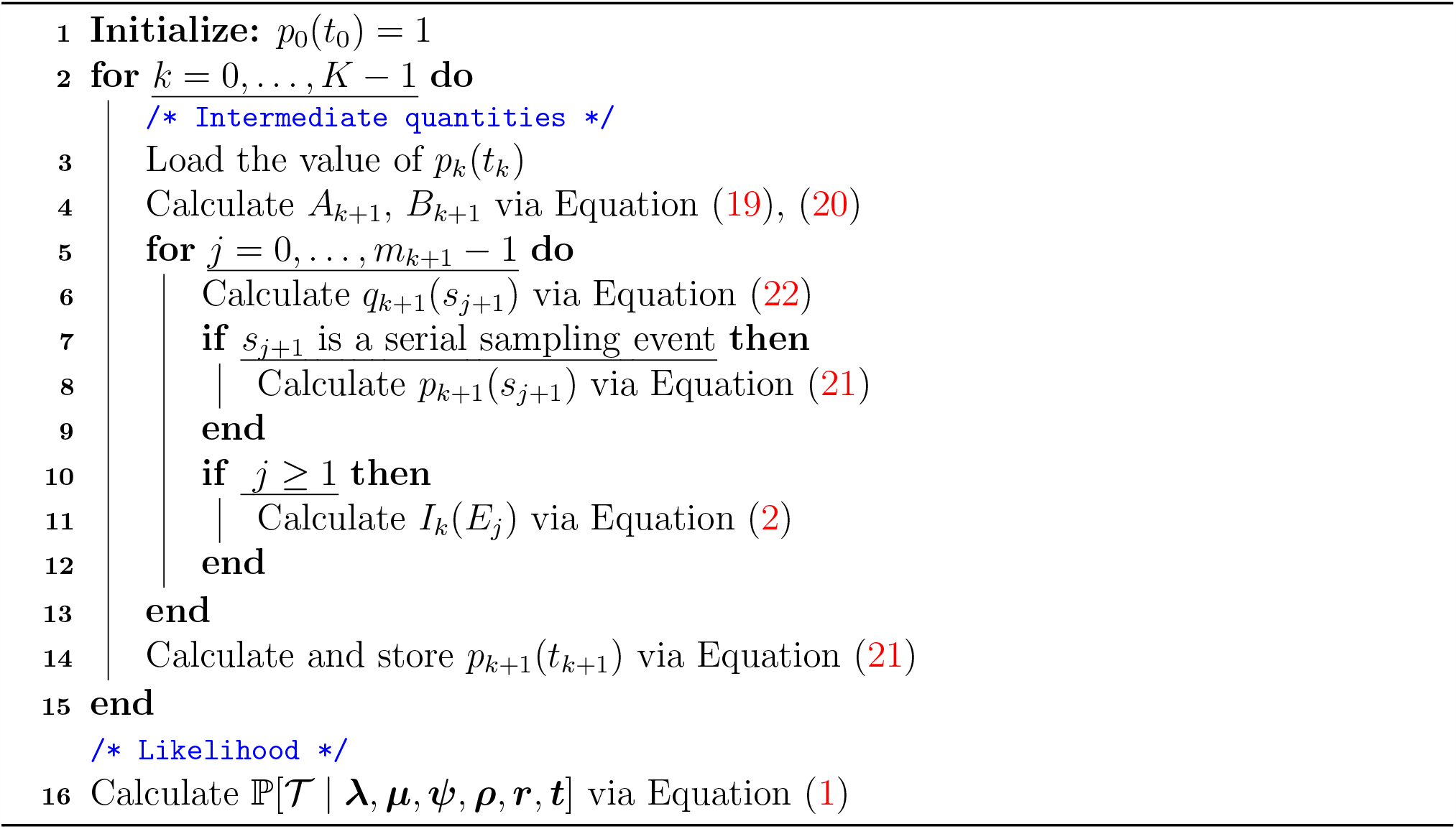

### S2 Gradient Derivation

#### S2.1 For 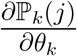

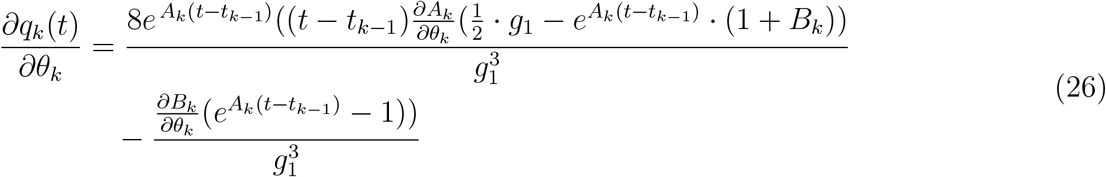

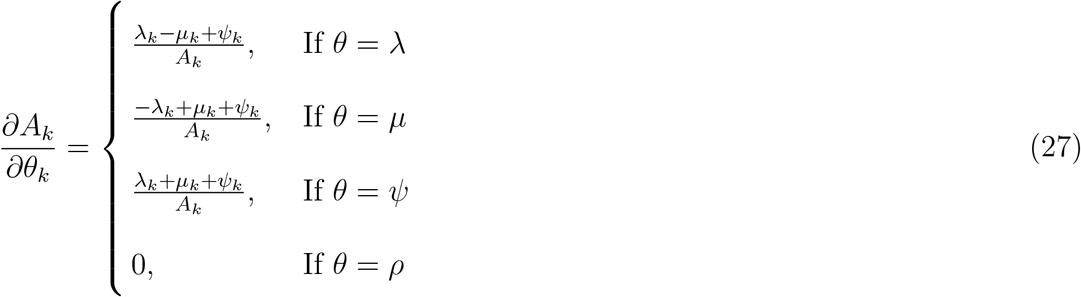

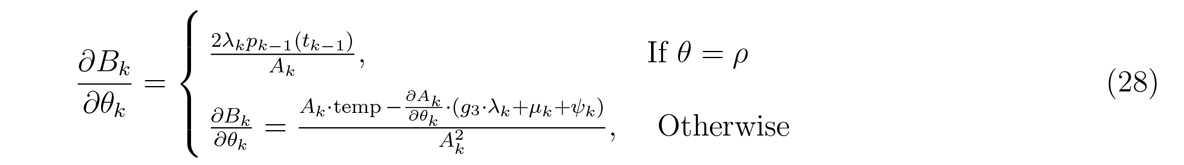

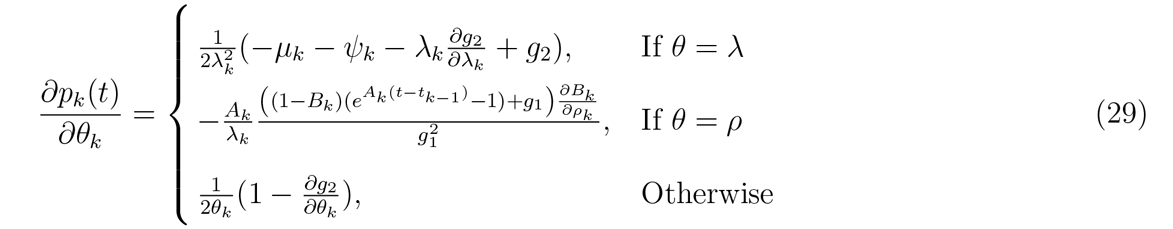

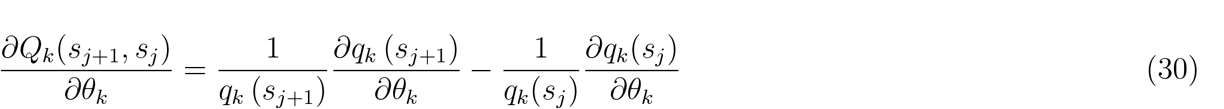

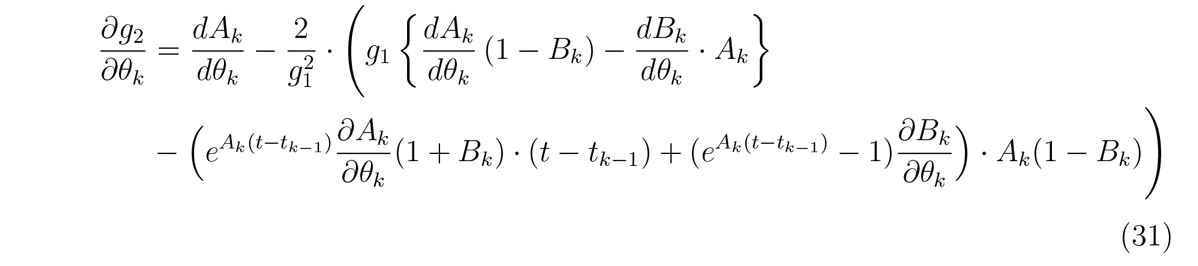

#### S2.2 For 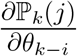 (*i* is an integer smaller than *k*)

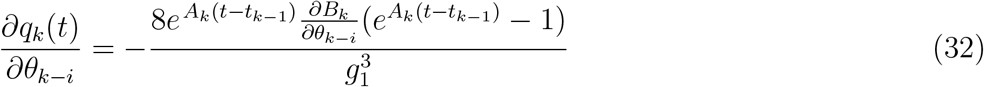

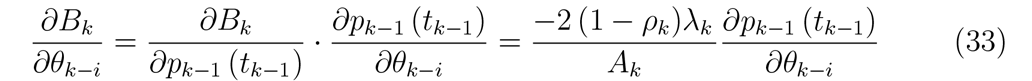

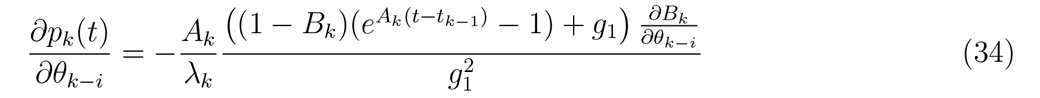

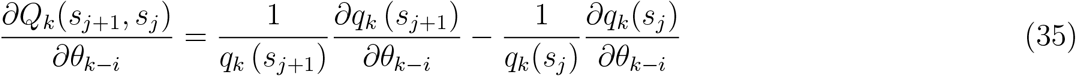

#### S2.3 Implementation Algorithm

We implement a recursive algorithm to compute the necessary gradient of the log-likelihood within our rate parameter space. Intermediate quantities are stored in between epochs to alleviate computational burden. Detailed algorithm is shown below based on the equations listed in 2.5 and previous sections in the supplement.

##### Algorithm 2

Gradient Calculation

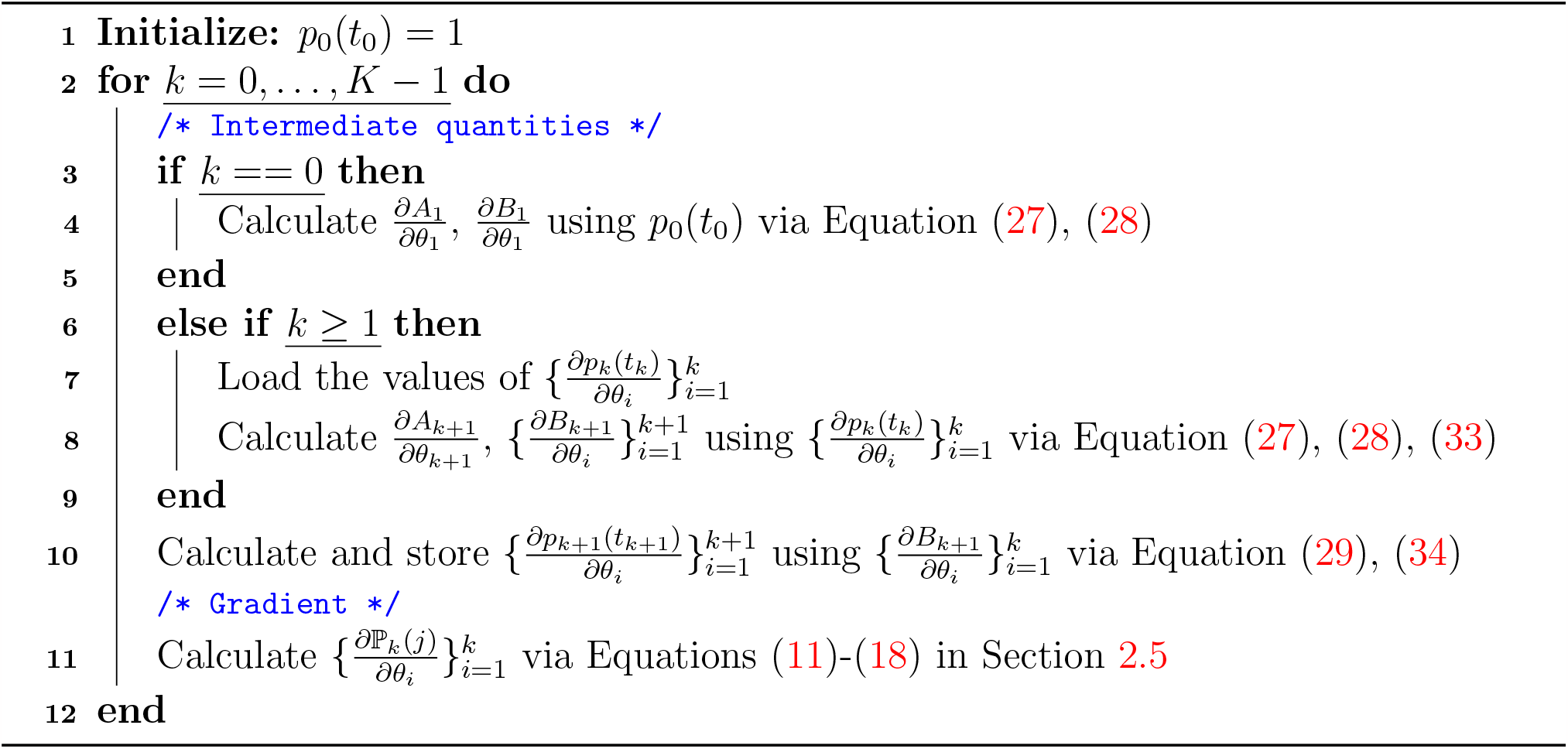

### S3 Prior distributions for EBDS models

#### S3.1 HIV dynamics in Odesa, Ukraine

We refer to the prior settings on the compound parameters from previous work (Vasylyeva et al. 2020), and try to roughly match their priors by adopting the following prior distributions on each of the rate parameters. Note that the sampling proportion was fixed to 0 before the first sampling date in their study, so we also set the sampling rate to 0 for the last two epochs for consistency.

**Table S1:** Prior specifications for the EBDS model in HIV virus analysis.

| Parameter | Prior | Role |
| --- | --- | --- |
| $\lambda$ | Lognormal (Mean = 0.85, SD = 1.0) | Birth rate |
| $\mu$ | Lognormal (Mean = -0.25, SD = 1.0) | Death rate |
| $\psi$ | Lognormal (Mean = -9.0, SD = 0.50) | Serial sampling rate |
| $t_{or}$ | Uniform (Lower = 19, Upper = 60) | Age of phylogeny |

#### S3.2 Seasonal Influenza in New York State

We follow the same framework for setting the priors for the GMRF-based model as in Section S3.3. Similarly, the prior distribution for the constant death rate is acquired by estimating the credible range for the duration of the infectious period according to reports by Centers for Disease Control and Prevention (n.d.), with 95% confidence intervals encompassing 6 to 11 days. Comprehensive information regarding the specific prior distributions is shown in the following table:

**Table S2:** Prior specifications for the EBDS model in Influenza virus analysis.

| Parameter | Prior | Role |
| --- | --- | --- |
| $\lambda_1^*$ | Normal (Mean = 3.08, SD = 1.17) | Log-scale birth rate at present |
| $\mu_k^*$ | Normal (Mean = 3.82, SD = 0.16) | Log-scale death rate for all epochs |
| $\psi_1^*$ | Normal (Mean = -0.77, SD = 1.17) | Log-scale sampling rate at present |
| $t_{or}$ | Normal (Mean = 12.5, SD = 15.0) | Age of phylogeny |
| $\alpha$ | Fixed to 2.0 | Exponent of the MRF |
| $\phi$ | Gamma (Shape = 1.0, Scale = 1.0) | Transformed global scale of the MRF |
| $\nu_k$ | Fixed to 1.0 | Local scale of MRF |

#### S3.3 Ebola epidemic in West Africa

We assume a constant death rate, *μ* for this data set, and we employ an empirical Bayes approach proposed by Magee et al. (2020) to set the prior on the first log-birth-rate and logsampling-rate in our Bayesian bridge MRF models. The prior for the constant death rate is obtained from an estimation of the plausible duration of infectious period with 95% confidence intervals covering 8 to 40 days (Velásquez et al. 2015). The detailed prior distributions can be found in the table below:

**Table S3:**
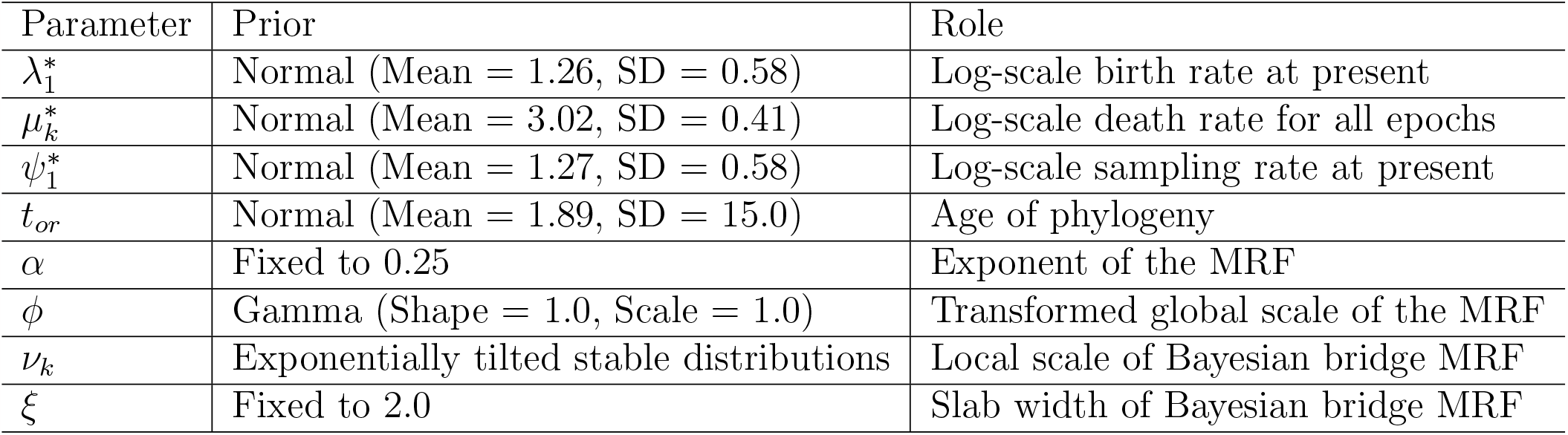
>Prior specifications for the EBDS model in Ebola virus analysis.

| Parameter | Prior | Role |
| --- | --- | --- |
| $\lambda_1^*$ | Normal (Mean = 1.26, SD = 0.58) | Log-scale birth rate at present |
| $\mu_k^*$ | Normal (Mean = 3.02, SD = 0.41) | Log-scale death rate for all epochs |
| $\psi_1^*$ | Normal (Mean = 1.27, SD = 0.58) | Log-scale sampling rate at present |
| $t_{or}$ | Normal (Mean = 1.89, SD = 15.0) | Age of phylogeny |
| $\alpha$ | Fixed to 0.25 | Exponent of the MRF |
| $\phi$ | Gamma (Shape = 1.0, Scale = 1.0) | Transformed global scale of the MRF |
| $\nu_k$ | Exponentially tilted stable distributions | Local scale of Bayesian bridge MRF |
| $\xi$ | Fixed to 2.0 | Slab width of Bayesian bridge MRF |

### S4 Inferred trajectories for birth/death/sampling rates

**Figure S1:**
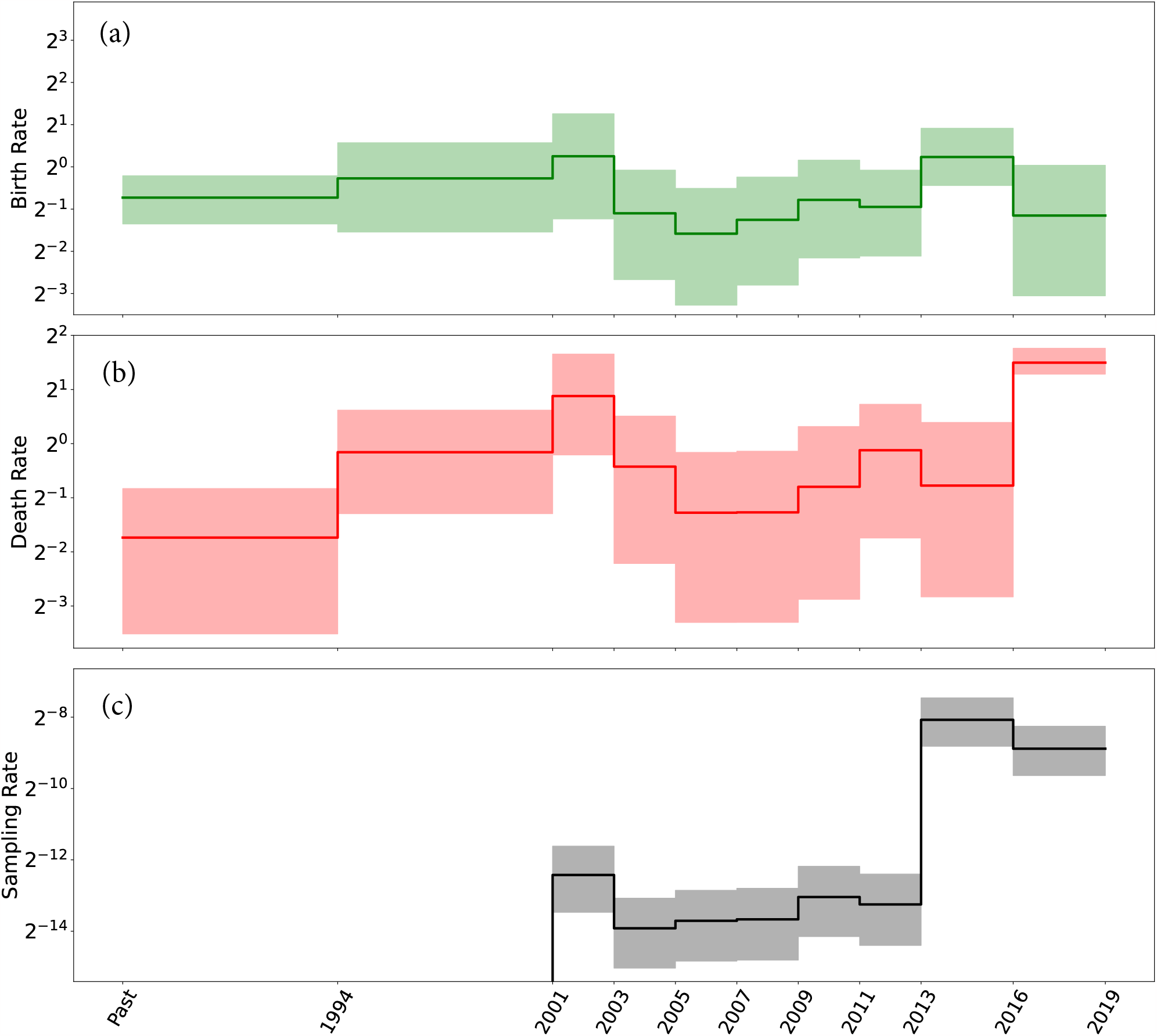
HIV virus: Median (solid line) and 95% credible intervals indicated by the shaded areas of the (a) birth rate, (b) death rate, and (c) sampling rate estimates through time.

**Figure S2:**
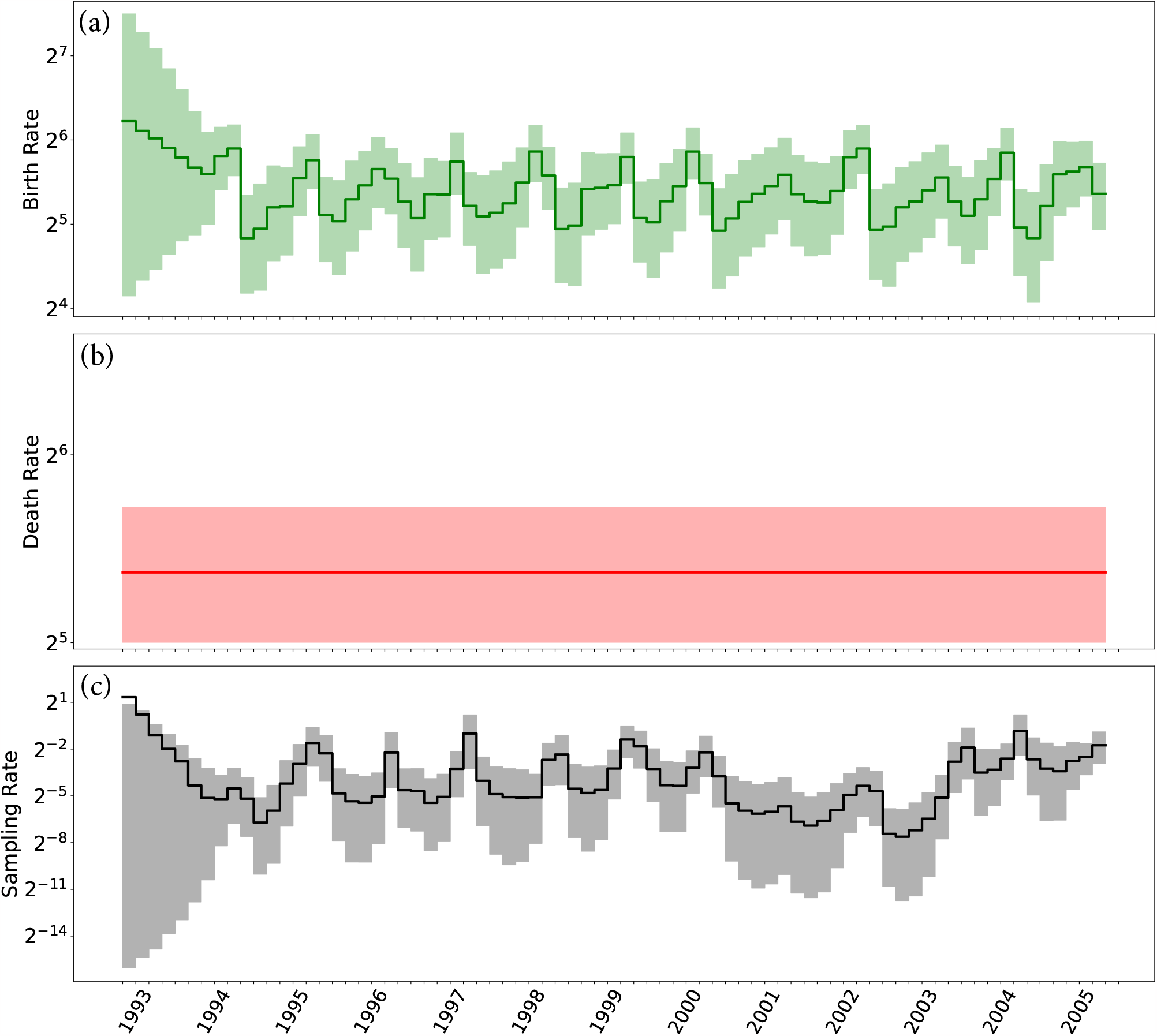
Influenza virus: Median (solid line) and 95% credible intervals indicated by the shaded areas of the (a) birth rate, (b) death rate, and (c) sampling rate estimates through time.

**Figure S3:**
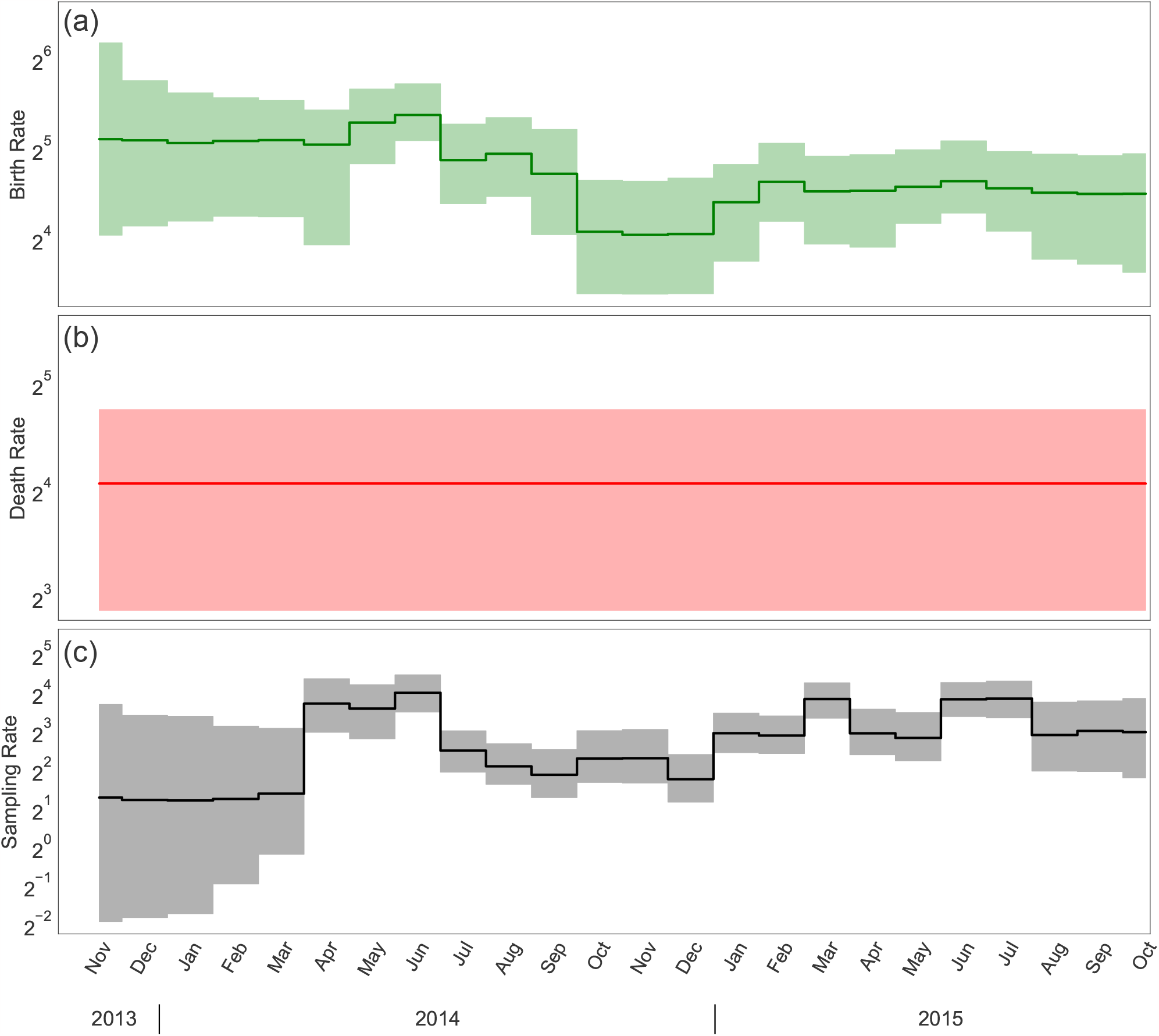
Ebola virus: Median (solid line) and 95% credible intervals indicated by the shaded areas of the (a) birth rate, (b) death rate, and (c) sampling rate estimates through time.

### S5 Computational complexity of the nodewise likelihood

The computational complexity of evaluating node-based representations of the likelihood is much less explicit. First, we need to write out an equivalent expression for the likelihood of Equation 1 node-wise. It will be helpful to distinguish different types of samples. In particular, let us denote serially-sampled tips ***ū***_*ψ*_ with a particular serially-sampled tip being *ū*_*ψi*_. With a slight abuse of notation, let us denote intensively-sampled tips ***ū***_*ρ*_, with ***ū***_*ρi*_ denoting the *vector* of intensively-sampled tips at the *i*th intensive-sampling event. Then we can write

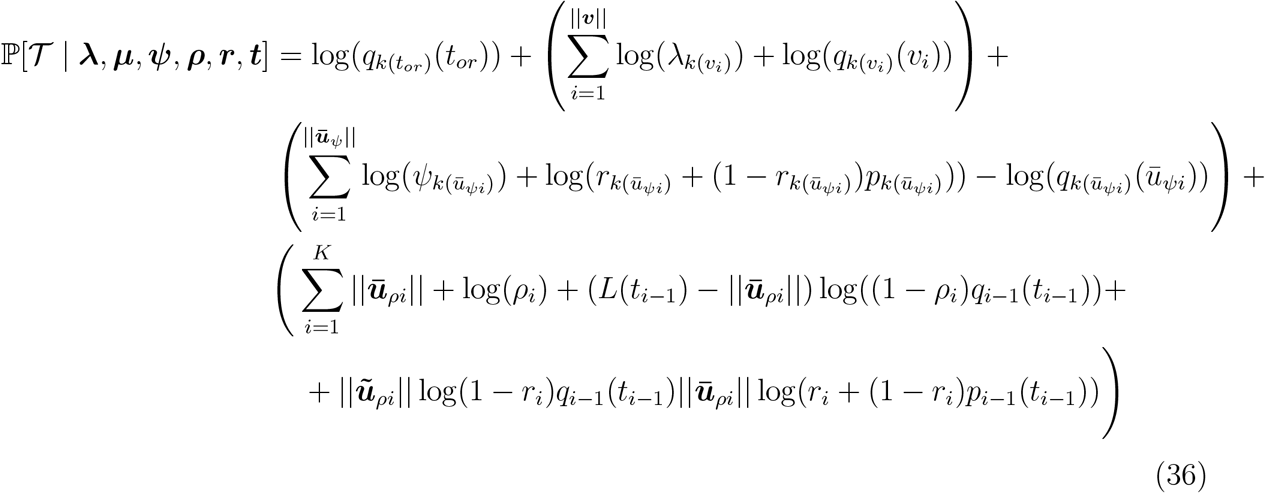

The complexity here is not immediately apparent for a number of reasons. For one, the complexity appears to depend on the relative proportion of samples of different types, which affects the number of values of *p*_*k*_(*t*) and *q*_*k*_(*t*) which must be computed. Importantly, the complexity of computing those *p*_*k*_(*t*) and *q*_*k*_(*t*) is not immediately apparent either, and that these costs are somewhat hard to disentangle, as *p*_*k*_(*t*_*i*_) builds recursively on *p*_*k*−1_(*t*_*i*_) and *q*_*k*_(*t*) depends on *p*_*k*_(*t*).

#### S5.1 Node lookups

Regardless of such ambiguities, all nodes in the tree require an interval lookup. For births, the lookup is required to find the correct *λ*_*k*_ term to use. For samples, the lookup is either to find the appropriate sampling rate, for serial samples, or to determine to which intensive-sampling event a sample belongs, for intensive samples. The time requirement here depends on the algorithm, for a binary search it is 𝒪 (log(*K*)), making the total lookup cost 𝒪 (*N* log(*K*)).

#### S5.2 How many computations of *q*_*k*_(*t*) are required?

In the worst, but most common, case, there are no intensive-sampling events and *q*_*k*_(*t*) must be computed for the times of all samples, all births, and all epoch times (note that even when *ρ*_*i*_ is 0, there is a term *L*(*t*_*i*_) log(*q*_*i*−1_(*t*_*i*_)) which must be computed in the final summation). In the best case, all samples are at intensive-sampling events, and *q*_*k*_(*t*) only needs to be computed for the times of all births and all epoch times. These are both 𝒪(*N* + *K*), though there is a factor of two’s worth of variation in front of the *N* depending on which side of this spectrum a tree falls in. Calling the cost of computing *q*_*k*_(*t*) *Q*, this makes the contribution to the complexity here 𝒪(*Q*(*N* + *K*)).

#### S5.3 How many computations of *p*_*k*_(*t*) are required?

The likelihood contains a number of explicit computations of *p*_*k*_(*t*) in the terms pertaining to (both serially- and intensively-)sampled tips. When all samples are serial samples, there are 𝒪 (*N*) direct computations of *p*_*k*_(*t*), while when all samples are intensive samples, there are 𝒪 (*K*). Taking the cost of computing *p*_*k*_(*t*) to be *P*, the addition to the cost here is between 𝒪 (*PN*) and 𝒪 (*PK*).

#### S5.4 What is the cost of computing *p*_*k*_(*t*) and *q*_*k*_(*t*)?

We have thus far shown that the cost of computing the nodewise likelihood appears to be between 𝒪(*N* log(*K*) + *Q*(*N* + *K*) + *PN*) and 𝒪(*N* log(*K*) + *Q*(*N* + *K*) + *PK*). But this is not particularly revealing without considering *P* and *Q*.

While *q*_*k*_(*t*) depends on *p*_*l*:*l<k*_(*t*) through **A** and **B**, once *A*_*k*_ and *B*_*k*_ have been computed, let us assume (as we did when evaluating the cost of the interval-wise likelihood) that the cost of *q*_*k*_(*t*) is 𝒪 (1). In other words, let us assume that 𝒪 (*Q*(*N* +*K*)) = 𝒪 (*P* (*N* +*K*)). This makes the implied cost of the nodewise likelihood between 𝒪 (*N* log(*K*) + *P* (*N* + *K*) + *PN*) and 𝒪 (*N* log(*K*) + *P* (*N* + *K*) + *PK*), which both simplify to 𝒪 (*N* log(*K*) + *P* (*N* + *K*)). Näively, we might choose to compute *p*_*k*_(*t*) recursively every time we need it, which is 𝒪 (*K*^2^). In this case, the implied cost of the nodewise likelihood is 𝒪 (*N* log(*K*) + *NK* + *K*^2^)).

#### S5.5 Precomputing **A** and **B**

One can instead choose to pre-compute *A*_*k*_, *B*_*k*_, as once these are computed the cost to compute *p*_*k*_(*t*) and *q*_*k*_(*t*) becomes 𝒪(1). Working backwards from the present allows recomputation to be avoided. As we did when we approximated the cost of the interval-wise likelihood, we will take the cost of the update (computing (*A*_*k*_, *B*_*k*_) from (*A*_*k*−1_, *B*_*k*−1_)) to be 𝒪 (1). Thus, the cost of the precomputation is 𝒪(*K*). This puts the implied cost of computing the nodewise likelihood between 𝒪(*N* log(*K*) + *N* + *K*).

#### S5.6 Counting lineages at epoch times

Regardless of whether the model includes intensive-sampling (that is, regardless of whether ***ρ*** = 0), one must compute *L*(*t*_*i*_) for all epoch times. This can be solved essentially the same way as the subintervals are obtained, at a cost of 𝒪(*N* + *N* log(*N*)). Alternately, it can be obtained by counting the number of births and sampled tips older (or younger) than each epoch time, at a cost of 𝒪(*KN*). This makes the lower end of the computational cost once again a range, from 𝒪(*NK* + *N* log(*K*) + *N* + *K*) to 𝒪(*N* log(*K*) + *N* log(*N*) + *N* + *K*).

In practice, the constants in front of all the sorting and node-lookup terms appear to be so small as to be unnoticeable in real-world computation. We demonstrate this in our timing experiments in the next section. Thus, for all practical purposes, the likelihood appears to be 𝒪(*N* + *K*) regardless of representation, as long as one avoids recursive computation of *p*_*k*_(*t*).

### S6 Timing Experiments

With the reformulation of the likelihood and derivation of the analytical gradients, our method notably gains in speed, as we highlight in this section. For a comprehensive assessment, we compare our approach with four other specialized packages for EBDS model inference concerning likelihood calculations. These include the BDSKY (Stadler et al. 2013) package within BEAST2 (Bouckaert et al. 2019), TreePar (Stadler et al. 2013) package in R (R Core Team 2021) and RevBayes (Höhna et al. 2016). Furthermore, we present a benchmark comparing the gradient calculation efficiency of automatic differentiation implemented in VBSKY (Ki & Terhorst 2022)package using JAX library (Bradbury et al. 2018) isolated from the variational inference procedure against our algorithm based analytical gradients implemented in BEAST.

To assess the scalability of the aforementioned methods in terms of likelihood/gradient calculation, we simulated a set of trees under the EBDS model with increasing number of tips. To investigate the scalability of different methods wrt the number of sequences, we fix the number of epochs to 5 for both likelihood and gradient calculation.

Regarding scalability with respect to the number of epochs, we adjust the model by progressively increasing the number of epochs. To keep other variables constant, we maintain the tree topology and set the number of tips at 12 (in scenarios where *K >> N*, this allows us to negate the effect of *N* in 𝒪(*N* +*K*)) for likelihood computation. For gradient calculations, we set the number of tips to 8198 (to minimize the impact of *K*^2^ in 𝒪 (*NK* + *K*^2^)).

For methods that employ just-in-time (JIT) compilation, including BEAST, BEAST2 and VBSKY, we run a short MCMC chain or variational inference algorithm to compute likelihood or gradient across 100,000 iterations and take the average run time.

**Figure S4:**
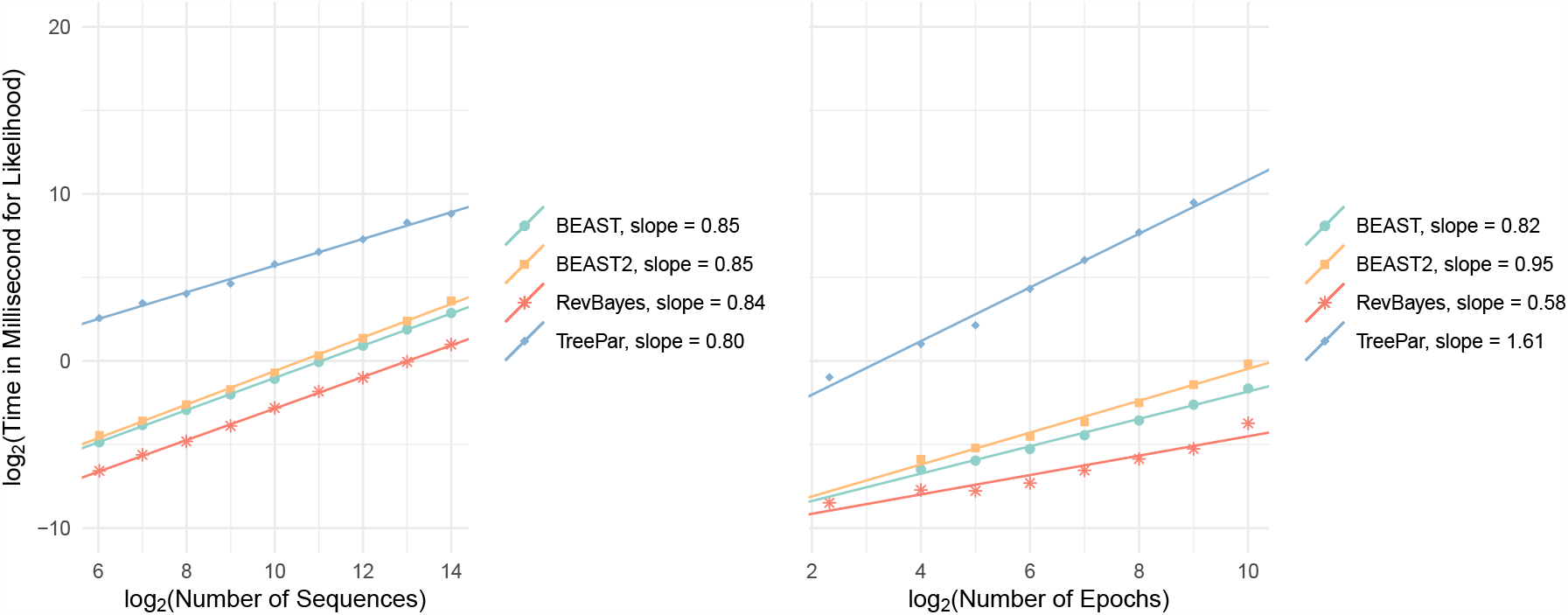
Speed of implementations for the likelihood calculations of increasing number of sequences (left plot) or number of epochs (right plot) for EBDS model. Note the time and number of sequences/epochs are laid out according to a logarithmic scale with base 2.

In our analysis, we observe that for likelihood computations, the implementations in BEAST, BEAST2, and RevBayes offer similar speed performance when adjusting both the number of sequences and epochs. In contrast, the TreePar package consistently lags, being several hundred times slower than its counterparts across all tested scenarios. It is also the sole implementation that exhibits a quadratic scaling with the number of epochs. The algorithms of BEAST, BEAST2, and RevBayes seem to demonstrate approximately linear scaling relative to both tree size and model epochs. It’s worth noting that RevBayes delivers the quickest calculation speed, which might be attributed to the inherent speed advantages of precompiled codes, particularly for quick likelihood calculations in our context. Result for TreePar with epochs exceeding than 512 is not not included as TreePar fail to process such large models.

**Figure S5:**
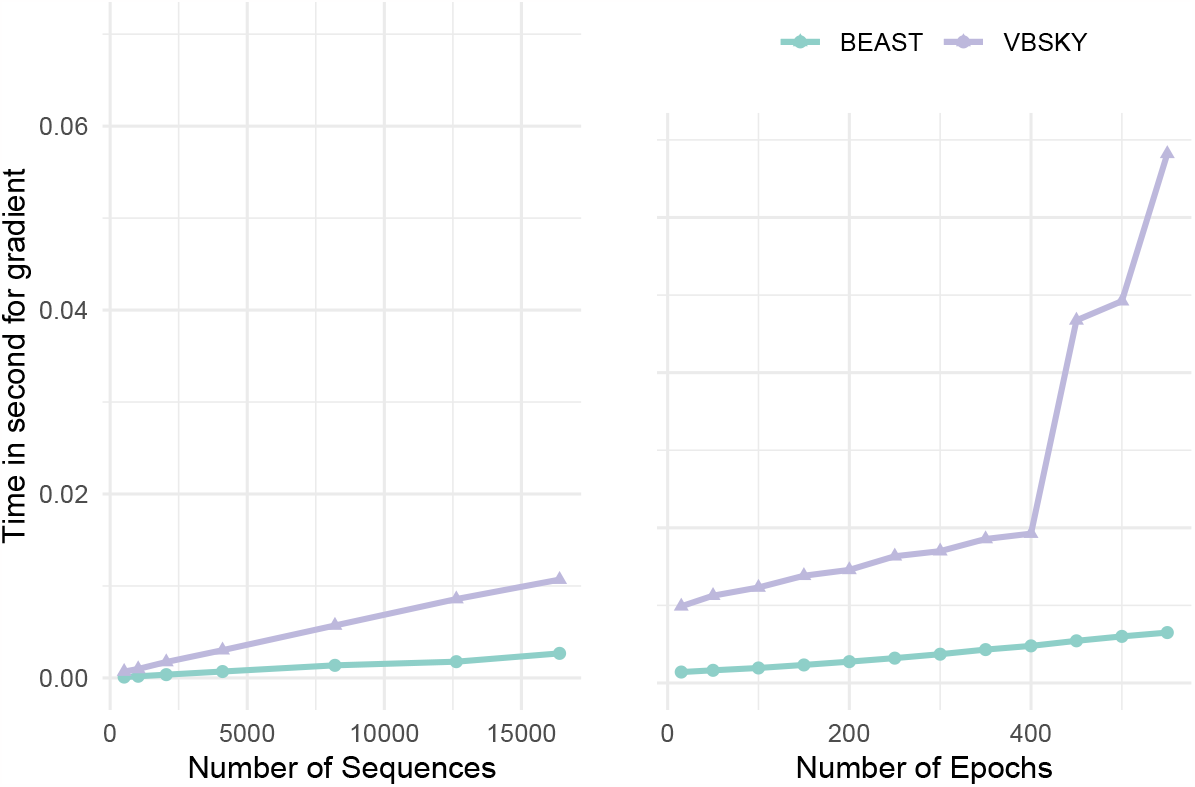
Speed of implementations for and gradient calculations of increasing number of sequences (left plot) or number of epochs (right plot) for EBDS model.

In terms of gradient calculations, our analytical gradients deployed within BEAST is remarkably faster than VBSKY approach using automatic differentiation. The gradient computation scales approximately linearly with the number of sequences for both BEAST and VBSKY. However, wrt the number of epochs, the scaling remains linear for BEAST but seems quadratic for VBSKY. We further confirm that the runtime slowness exhibited in VBSKY is not due to memory issues or JIT compilation difficulty. Therefore, our analysis demonstrates that analytically calculating the gradients of the EBDS likelihood is critical for improving the running time of gradient based methods.

